# Statistical structure of locomotion and its modulation by odors

**DOI:** 10.1101/404772

**Authors:** Liangyu Tao, Siddhi Ozarkar, Jeff Beck, Vikas Bhandawat

## Abstract

Most behaviors such as making tea are not stereotypical but have an obvious structure. However, analytical methods to objectively extract structure from non-stereotyped behaviors are immature. In this study, we analyze the locomotion of fruit flies and show that this non-stereotyped behavior is well-described by a Hierarchical Hidden Markov Model (HHMM). HHMM shows that a fly’s locomotion can be decomposed into a small number of locomotor features, and odors modulate locomotion by altering the time a fly spends performing different locomotor features. Importantly, although all flies in our dataset use the same set of locomotor features, individual flies vary considerably in how often they employ a given locomotor feature, and how this usage is modulated by odor. This variation is so large that the behavior of individual flies is best understood as being grouped into at least 3-5 distinct clusters, rather than variations around an average fly.

## Introduction

There are many approaches to the study of neural underpinnings of behavior: One large body of work is rooted in the psychophysical literature where an animal is forced to choose between a few discrete behaviors^1^. This approach allows one to control stimuli and rigorously analyze behavioral outcomes based on an established framework^2^, but sacrifices a full analysis of behavioral dynamics, leaving critical issues unexplored. Another large body of work is focused on understanding behaviors that are reflexive (albeit with some flexibility) such as saccades^3^, collision avoidance in insects^4^ and others. Another large body of work has focused on the control processes involved in goal-directed behaviors such as reaching movements and have revealed many fundamental principles of motor control. Yet, another popular behavioral motif that has received much attention is behaviors that require meticulous sequencing^5^. Finally, much work has been done to elucidate the workings of central pattern generators that underlie the rhythmic motor activity during walking and running^6^. Although many of the relatively stereotypical behavioral motifs described above are at play during most behaviors, they are not helpful in describing the structure underlying most everyday activities such as making a cup of coffee or a peanut-butter sandwich or walking to a car which consist of a sequence of actions, but neither the sequence nor each sub-action is stereotyped. These activities and their underlying sub-actions cannot be described either as sensorimotor reflexes or as behaviors that arise out of meticulous sequencing. An important example of such a behavior is an animal’s locomotion. While tracks of a mouse or a fly exploring a chamber are not stereotypical there is an obvious structure to it.

Uncovering the structure within non-stereotyped behaviors such as locomotion require sophisticated analytical tools which can be employed on two complementary representations of an animal’s behavior: In recent years much progress has been made in the use of analytical tools which describe behaviors as transformation in the shape/posture space using both supervised^7,8^ and unsupervised^9–12^ algorithms. These studies have provided remarkable insights by showing that much of an animal’s behavioral repertoire can be described as transitions between a small number of postures. While behavioral descriptions as transformations in posture space accurately classify the type of behavior (such as locomotion vs. grooming), because an animal’s position in the world coordinate system is ignored, the structure within an animal’s trajectory in world coordinates remains relatively unexplored.

Much of the work in extracting structure from an organism’s trajectory is derived from the “run and tumble model” which was originally employed in the context of bacterial chemotaxis^13^. In this model, the organism is assumed to travel in relatively straight lines (runs) of exponentially distributed run lengths until they make sharp turns (tumble) to choose another direction at random. It is tempting to consider the motion of larger animals as roughly approximated by a run and tumble model, and many studies (explicitly and implicitly) employ a run-and-tumble framework with increasing sophistication as an analytical framework for locomotion^14–16^. However, it is clear that run-and-tumble based model cannot describe an animal’s locomotion. One obvious well-documented limitation is that animals do not turn at discrete times^17–22^ and therefore cannot be described using a run-and-tumble framework. More generally, larger animals are likely to exert greater control over speed and direction of their locomotion and a better model is necessary to understand the resulting structure in its trajectory.

The lack of a model for locomotion makes it difficult to quantify the effect of stimuli on locomotion and is a critical missing piece in understanding sensorimotor transformations. As an example, in most studies of odor-modulation of locomotion, odors are primarily described as attractive or repulsive; this description is based on the end result, and does not consider the navigational maneuvers that underlie these end-results. Ignoring the underlying navigational maneuvers have led to a fundamental misunderstanding of odor modulation of locomotion. In a recent study, based on detailed analysis of a fly’s locomotion, we have demonstrated that the navigational maneuvers in response to similarly attractive odors are quite distinct^20^. The analysis used in that study was based on an ad-hoc parametrization of locomotion, and not on a generative model of locomotion, making it difficult to determine whether our chosen parameter set was appropriate. A model of locomotion also makes it possible to compare how locomotion is affected by a given stimuli, and also how different individuals differ in their locomotion and in their response to stimuli.

In this study, we employ a hierarchical statistical model, Hierarchical Hidden Markov Model (HHMM) to describe the structure in the fly’s locomotion^23^. We show that a fly’s locomotion is well-structured and that HHMM is an elegant representation of this structure. HHMM provides a simple and intuitive description of both a fly’s locomotion and the effect of odors on the same. Surprisingly, different flies employ different strategies in their locomotion both before odor onset and in their response to odors. Our data is, thus, inconsistent with the idea that the behavior of different flies represents variations around an “average” fly. Rather, our data is most consistent with the idea that flies employ 3-4 different strategies, at a minimum, to explore a small circular arena and a similar number in their response to odors.

## Results

### Rationale for the choice of HHMM as the model and the model architecture

We model the locomotion of wild-type flies exploring a circular arena^20^ whose center (odor-zone) consists of a fixed concentration of odor (Figure 1A). The arena and the experimental procedure is detailed in an earlier manuscript^20^. Briefly, locomotion of each of the 34 flies in our dataset was measured 3 minutes before an odor (apple cider vinegar or ACV) was turned on, and 3 minutes during the presence of ACV. Sample trajectories are shown in Figure 1B.

**Figure 1.**
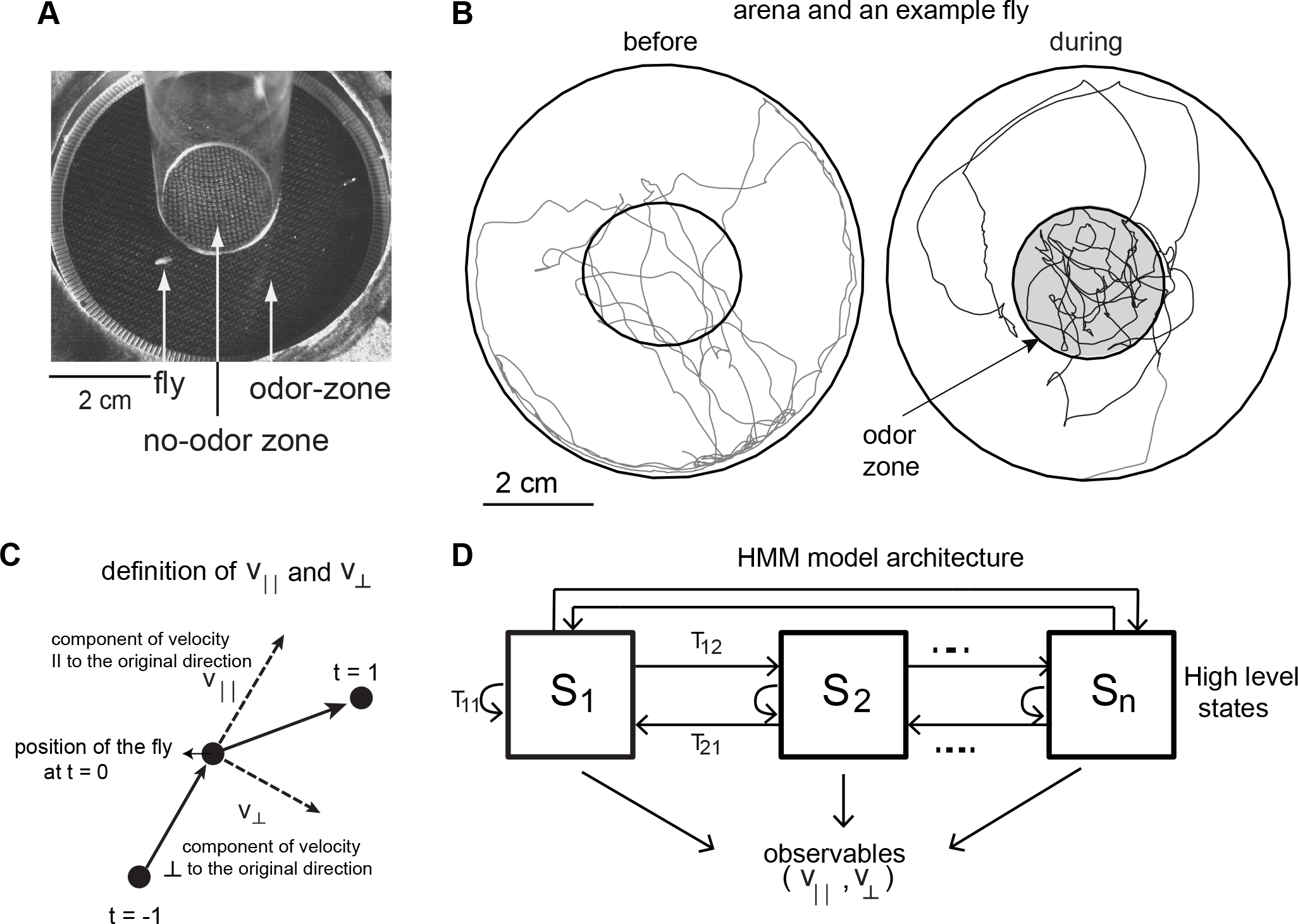
Experimental Setup and HMM architecture. **A.** Top view of the chamber. **B.**Tracks of an example fly in a circular arena (3.2 cm in radius). The central region (1.2 cm) has no odor during the 3 minute period before the odor (before period) and is odorized in the “during” period. The odor zone is shaded. **C.**The observables - V_││_ and V_⊥_ at time point t=0 is schematized. V_││_ is the component of the fly’s velocity along the velocity vector at the previous time point; V_⊥_ is the component perpendicular to the velocity vector at the previous time point. **D.**HMM Model Architecture: A single layered model which has n number of states which are defined by a joint probablity distribution of the observables. The probability of transitioning from the *i*^*th*^ HL state to the *j*^*th*^ HL state is given by Tij.

Inspired by past success at modeling animal trajectories using Hidden Markov Model (HMM), we attempted to model the fly’s locomotion using a HMM^24 25^. HMM creates discrete states based on a time series of observables such as position, speed or acceleration. The advantage of using HMMs in modeling locomotion is well described in earlier studies^24^. Briefly, instantaneous measures of an observable are variable; therefore, behavioral states inferred by simple thresholding applied to instantaneous measures of the observables are likely to be more erroneous. HMM remedies this problem by inferring states based not only on the value of the observable at the current time point but also on the previous and following time points and allows a more accurate determination of state (this idea is well-explained in Figure 2 in ref 17).

**Figure 2.**
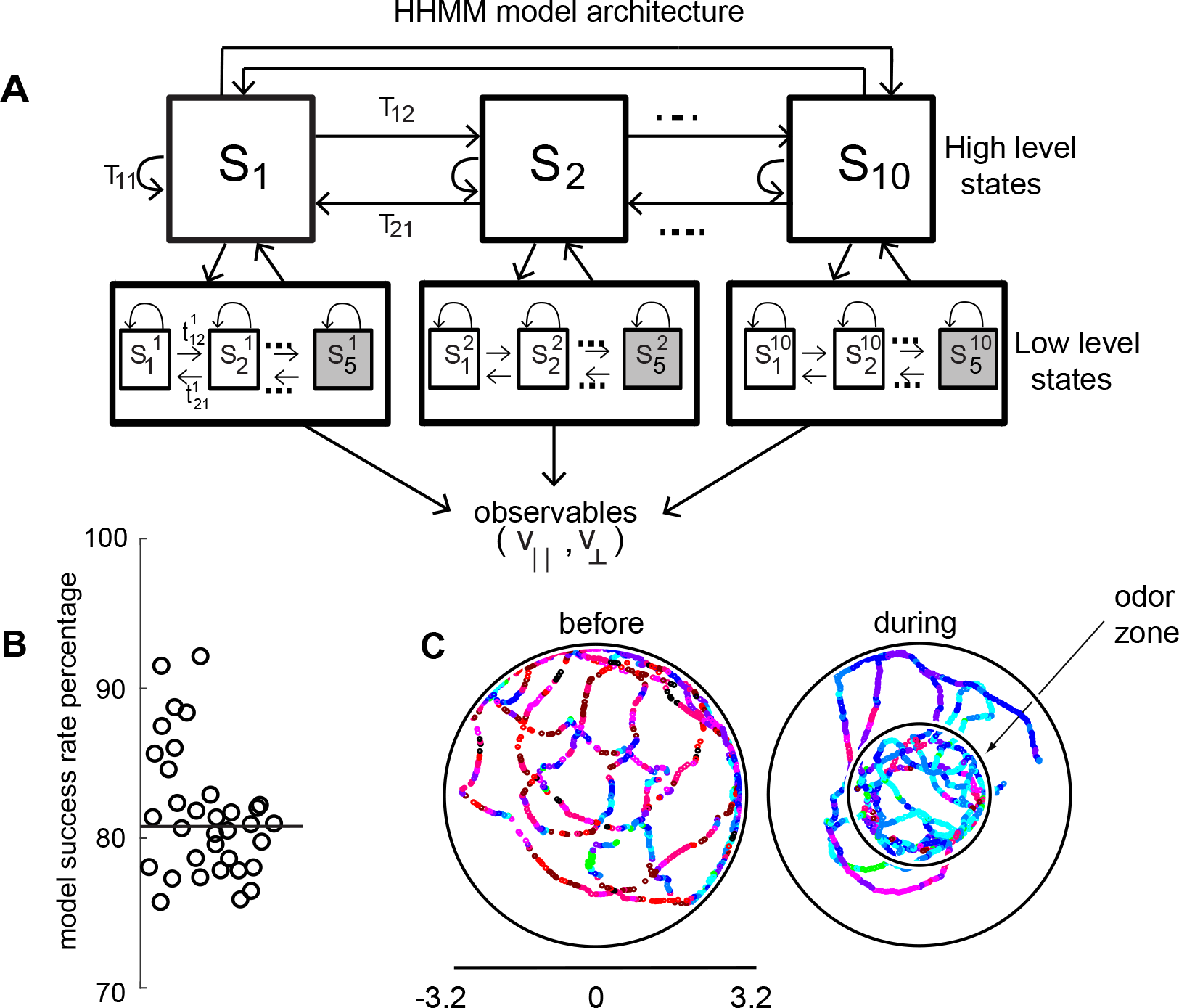
HHMM Model architecture. **A.** Model architecture: The model consists of two layers; there are 10 high level states (HL states) each of which have 5 low level (LL) states. The probability of transi-tioning from the *i*^*th*^ HL state to the *j*^*th*^ HL state is given by Tij. At the lower level, each HL state has its own transition probability matrix that describes transitions between its LL states. The shaded boxes represent the terminal states. **B**. As a measure of the model’s ability to fit the data for individual flies in our dataset-the percentage of timepoints for which the model had >85% confidence is plotted. **C**. HL state assign-ment for a single fly. The 10 high level states,each coded using a different color are overlayed on the tracks of a fly that were observed before(left) and during(right) the odor stimulus period.

In this study, we use observables that describe the change of position as a function of time, and hence our analysis will focus on behavioral states in the velocity space. The most commonly used representation of locomotion in the velocity space is instantaneous speed and angular speed. But, as noted by others^24^, because it is difficult to measure angular speed accurately at low speeds (see methods for details), we fit the model to two other measures of speed - the components of the fly’s speed, at each time point, parallel 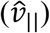 and perpendicular 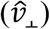 to its movement during the previous time point (Figure 1C, Methods section 2). If the fly walks straight ahead, 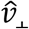 would be zero, therefore, 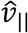 and 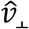 are closely related to speed and angular speed. We fit a time series of 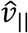 and 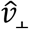 to an HMM. The HMM architecture is shown in Figure 1D. We employed models with 24 to 50 states. HMM was only modestly successful as a model. One evidence for this modest performance is that HMM was never able to classify >70% of the tracks into one of the states with >85% confidence. We will return to why HMMs perform modestly when we compare its performance to a HHMM.

The transition probability matrix for the HMM was sparse (Figure 1-S1) suggesting that from each state there are transitions to only a handful of other states. One method to improve upon the performance of HMM is to cluster the states obtained by HMM according to the transition probability matrix. A common approach to clustering is to block-diagonalize the transition probability matrix^9^. Tracks corresponding to the 10 states obtained by clustering are shown in Figure 1-S1. Some of these states appear to describe recognizable features in the data such as left (state 10) or right turn (state 9). But efforts to block-diagonalize the transition probability matrix were only partially successful. The most obvious failure corresponds to the states with little movement. These states – describing the absence of movement – can occur in many different contexts, such as the fully stopped state or intermittent runs. When the same state is used in different contexts, an approximate block diagonalization of the transition probability matrix would fail because the state would belong to two blocks of the block-diagonal. In these cases, the different states that correspond to the absence of movement are not clustered at all and appear alone even after block-diagonalization. Thus, the existence of the same state in different contexts is one important reason for the modest success of HMM, this limitation of HMM is well-known^26–31^. Another reason that an HMM is modestly successful is that the duration of states modeled by HMM are shorter than the states in the data (this point is expanded later). In essence, HMM is only moderately successful because states with similar distribution over observables occur in different contexts, and because HMMs create states with shorter duration.

Because of the modest success of the HMM and an approximate block diagonal structure of its transition probability matrix, we evaluated the ability of a 2-layered Hierarchical Hidden Markov model (HHMM) to model the data (Figure 2A). We reasoned that the low-level states (LL states) would be represented by Gaussian distributions on the observables, and the high-level states (HL states) would therefore be a mixture of Gaussians and would be better able to model the experimental data. Moreover, these HL states would have longer duration than the states discovered by HMM allowing it to more naturally model composite states.

Indeed, in a Bayesian model comparison (see methods), HHMMs outperformed HMMs with the same number of states. Since HMMs rarely used more than 35 states (even when models with higher number of states were fit), to perform model comparisons, we used models with less number of states than the particular HHMM we will eventually employ. We compared a 2-level HHMM with 6 HL states and 4 LL states to a single-level HMM with the same number of states (4 × 6 = 24 states). If we label the non-hierarchical model as the null model, we were able to reject the null model using Bayesian model comparison at p<0.0001, implying that a hierarchical model is necessary. Another model comparison – a 2-level model with 8 HL and 4 LL states compared to a single-level model with 32 states – also yielded similar results. The objectively better performance of a HHMM compared to a HMM suggests that a model that includes a hierarchical structure is more consistent with a fly’s locomotion. It is important to note that, despite its name, the HHMM is actually a simpler model compared to a HMM with the same number of states. This is because any HHMM – which puts very specific constraints on the transition probability matrix – can be represented by an HMM but not vice-versa. In other words, HHMM with the same number of states has far fewer parameters (see methods section 4). Thus, for the two comparisons above, HHMM has 6^2^ + 6*4^2^+6*4*5 = 252, and 8^2^ + 8*4^2^ +8*4*5 = 352 parameters, and HMM has 24^2^+24*5 = 696 and 32^2^+32*5 = 1184 parameters respectively; therefore HHMM has fewer parameters. When a simpler model provides a better characterization of the data, then we can conclude that the additional structure contained in that model provides a more accurate characterization of the structure in the data.

The fitting procedure and a detailed description of the model is in the methods (section 3). The model we chose has 10 HL states (Figure 2A) and 5 LL states for each HL state. The model was fit to the entire dataset – both before and during the presence of odors. The fitting process initializes by fitting each fly’s tracks to its own HHMM and then clusters these 34 HHMMs – one for each fly – resulting in a smaller number of models. Remarkably, a single HHMM is an excellent fit for all the data suggesting that the behavior of wild-type are composed of similar components. One evidence that a single model is an excellent fit is that the model was able to successfully assign an HL state (defined as >85% confidence) for >80% of the data points (Figure 2B, median 81%). This percentage was consistently high for all flies in our dataset (Figure 2B). In comparison, an HMM with 50 states can only classify 68% of the data with the same level of confidence. Tracks of a fly with each HL state labeled with a different color are shown in Figure 2C.

As noted above, the greater success of HHMM derives from its ability to distinguish between states with overlapping distributions based on the transition probability of these states, and the longer duration of its states. To make this clearer, we compare the HHMM above to a HMM. First, as expected the time a fly spends in a HMM state is shorter than the time it spends in a HHMM state (Figure 2-S1A). The longer time a fly spends in a HHMM state is a result of the hierarchical structure, and allows a HHMM to more accurately assign states, and is illustrated with two examples. First, consider a track that is assigned as a left turn by HHMM, the HMM only classifies parts of the track as a left turn because of the inability of HMMs to consider longer duration trends in the observables (Figure 2-S1B). Short-term inhomogeneity in the data throws the HMM off; as soon as the 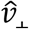 decreases, the HMM exits its left turn state. Another example, this time for the nearly stopped state, is shown in Figure 2 – S1C. In this example, too, the HMM exits the stopped state as soon as there is a small movement. The net result is that the HHMM is able to classify all long stops into a single state while HMM needs four different stop states. HHMM also assigns more of the left turn as such. The net result is that HMM assigns only 2% of tracks as left turn compared to 6% assigned by the HHMM. HMM also has four stop states with nearly the same observable distributions.

The limitations of HMM in describing phenomenon which have hierarchical and shared structure because of the short duration of its states is well documented^26–31^. In our case, we show that HHMM is objectively a better model of data than is a HMM.

### HHMM reveals that a fly’s locomotion is surprisingly structured in the velocity space

The HHMM we present captures how a fly’s locomotion is structured in the 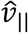 and 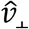 space essentially the change in a fly’s position as a function of time. This structure is evident in both the organized transitions between HL states and the narrow range of observables associated with each state: The organized transitions between HL states are shown in the transition probability matrix in Figure 2-S2. The transition probability matrix is sparse – a vast majority of transitions from each HL state were to 2-3 other HL states. When we reordered the states (see methods section 5) from low-speed-high-turn-states to high-speed-low-turn states, we found that from any state the flies transitioned to the neighboring states with a high probability (Figure2-S2) suggesting a gradual transition from low-speed-high-turn states to high-speed-low-turn states.

This gradual transition is not because flies cannot make large transitions due to biomechanical limitations because 47 out of the 81 possible transitions between HL states have a non-zero probability. Rather, transitions to states with similar kinematics show that under our experimental conditions – locomotion in a dark, small circular arena - flies locomote at similar 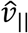 and 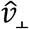 for extended time periods and represent one way in which locomotion is organized.

More important to the organization is the narrow distribution of observables - 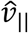 and 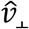-associated with each HL-state. The structure of one HL state (state 10) is shown in Figure 3. The distribution of observables for the HL state is a composite of the distributions of its LL states (Figure 3A). Both the model (solid line in Figure 3A1) and a random sample of observables drawn from the time points assigned to a given LL state (gray markers) show that during each LL state within HL state 10, the observables are limited to a narrow range of values; in each LL state belonging to HL state 10, 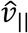 is large and 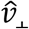 is negative implying that in HL state 10, flies turn counter-clockwise at high speeds as seen in sample tracks corresponding to a single transition to HL state 10 (Figure 3B). Fast, counter-clockwise turns represent a *locomotor feature* which describes a fly’s locomotion in HL state 10. To better visualize this feature, we translated each track such that it began at the origin, and rotated the tracks so that the initial velocity vector pointed along the y-axis (Figure 3C1, see methods). These transformations make it apparent that the locomotor feature for state 10 is turning left at high speeds (Figure 3C2).

**Figure 3.**
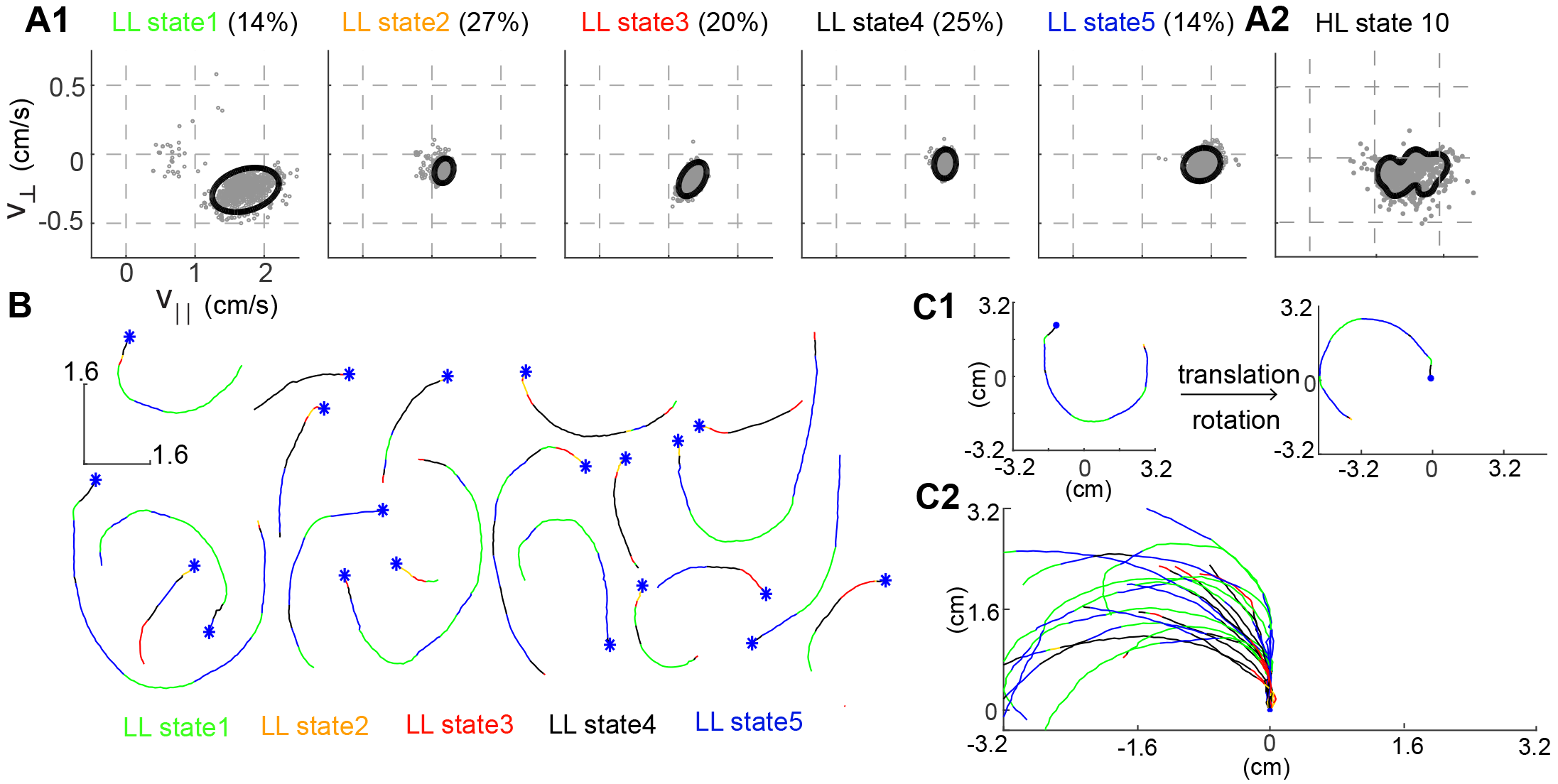
Structure of a HL state. **A1**. 85% confidence bounds of the model (black ellipse) and randomly selected observables (gray dots) corresponding to data points assigned to the LL states underlying HL state 10. Percentage of time spent in a given LL state is also shown. **A2.** Distribution of observables for the HL state. **B.** Example tracks denoting a single transition to HL state 10 show that the fly is turning counterclockwise. LL states were color coded. **C1.** Each track is rotated and translated for visualization. **C2.** All 20 tracks in B were transformed as shown in C1. Transformation reveals that all HL state 10 trajectories represent left turns.

Just as HL state 10 has a narrow distribution of observables and corresponds to a locomotor feature, the other HL states also correspond to distinct locomotor features. To visualize the locomotor feature represented by each HL state, we rotated and translated (as in Figure 3C) tracks corresponding to each of the 10 HL states (Figure 4). The distribution of the observables – 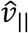 and 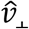 - for each HL state is also plotted. HL state 1 represents very slow walking with frequent changes in direction. In state 2, flies are either completely stopped or they walk at a speed about twice the speed of the fly in state 1; state 2 represents stop and start locomotion. The subtle, but important differences between state 1 and 2 show an instance in which the HHMM is successful at extracting an unexpected feature in the velocity profile in a fly’s locomotion. Moreover, when it does walk, it walks at speeds greater than the speed in state 1. During state 3, the fly is exhibiting a sharp turn. This sharp turn is reflected in the increase in 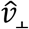 with a concomitant decrease in 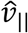. These three states together represent slow locomotion.

**Figure 4.**
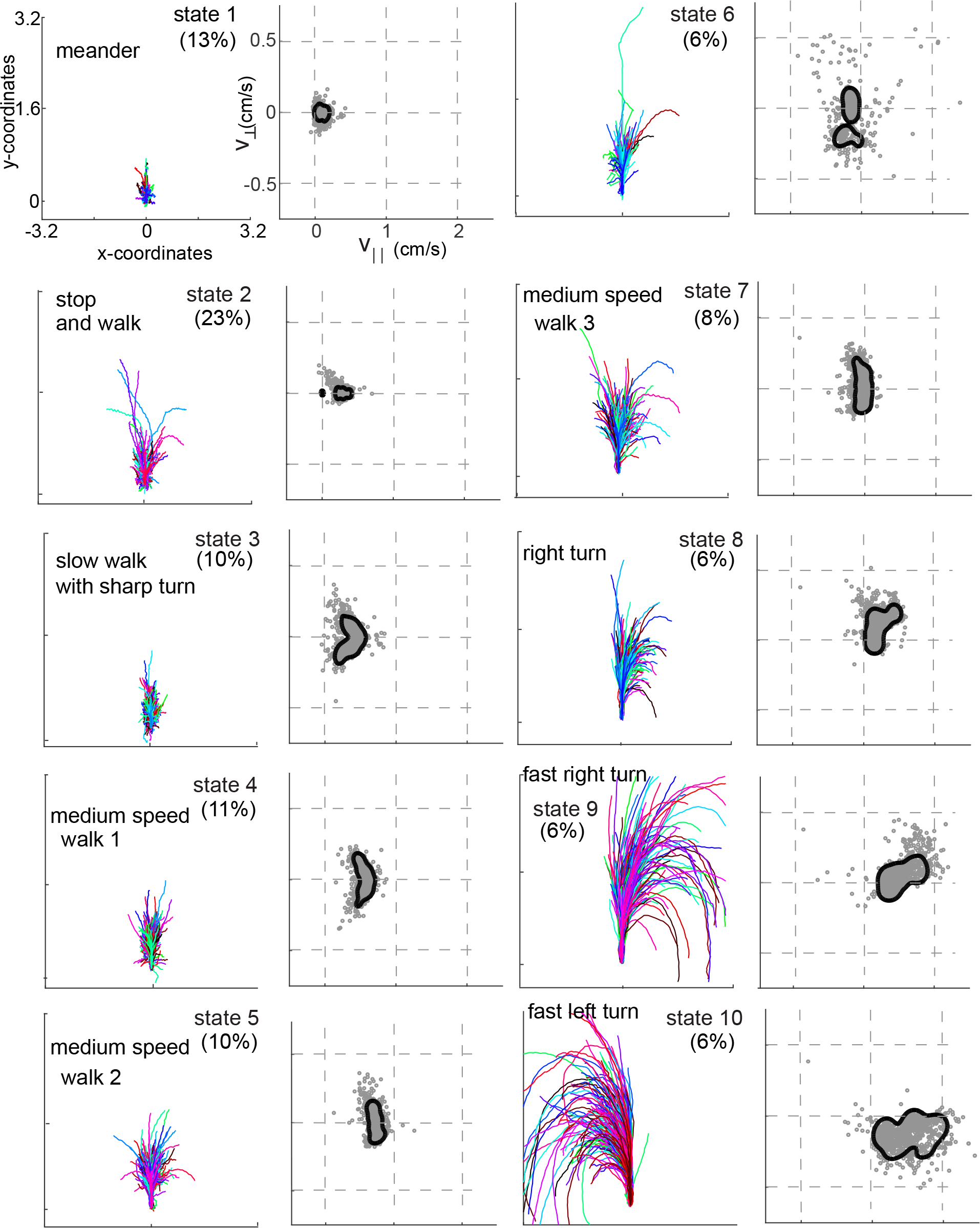
Each HL state describes a locomotor feature during which the fly uses a narrow region of the velocity space. Trajectories (left) and distribution of observables (right) corresponding to each of the 10 HL states are shown. Trajectories were transformed via translation and rotation to start at the origin and moving straight up. Both model distribution (solid lines) and randomly selected 1000 empirical datapoints (gray dots) belonging to a given state are shown. The percentage of occurence of each HL state is given.

In states 4-7, flies are walking at a medium-speed. In contrast to the clear drop in 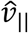 with increases in 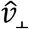 in state 3, 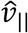 remains strikingly constant irrespective of 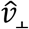. These states are different from each other because 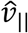 is slightly different.

States 8-10 are high-speed states; each of these states is also characterized by their turn direction. During states 8 and 9, the fly turns right; the fly’s speed is higher during state 9 than during state 8. During state 10, the fly turns left. States 9 and 10 are mirror-symmetric versions of each other.

In sum, HHMM performs better than a HMM at modeling a fly’s locomotion. As noted by others^24^, locomotor states cannot be reliably determined by looking at a single time point. HHMs allow more reliable state determination by looking at adjacent time points. HHMM further extends the benefits of a HMM by lengthening this temporal window (as illustrated in Figure 2-S1). In our dataset, HHMM discovers locomotor features – defined by periods when the flies occupy a characteristic, small region in the velocity space - that last over extended periods of time: they spend 60% of time performing a locomotor feature for >300 milliseconds, and >10% of their time performing a single locomotor feature for >3s (Figure 2-S1). Remarkably, because the same HHMM fits the track of all the 34 flies in our dataset, it allows us to describe locomotion and the effect of stimulus on locomotion based on these 10 locomotor features.

### Odors affect locomotion by altering the occupancy of HL states

We considered two general possibilities for modulation of locomotion by odors: First, since the HHMM was fit to the entire dataset – both before and during the presence of odor - it is possible that some of the 10 HL states are employed exclusively in the presence of odor. Alternatively, it is possible that although all the HL states are employed even before the introduction of odor, the frequency with which they are used changes in the presence of odor. To distinguish between these possibilities, we measured HL state occupancy before odor onset and during the presence of odors. Because our previous study with the same dataset^20^ had shown that even though the odor is present only within the odor-zone, it affects a fly’s behavior both inside and outside the odor-zone, we separately investigated changes in state occupancy inside and outside the odor-zone (Figure 5).

**Figure 5.**
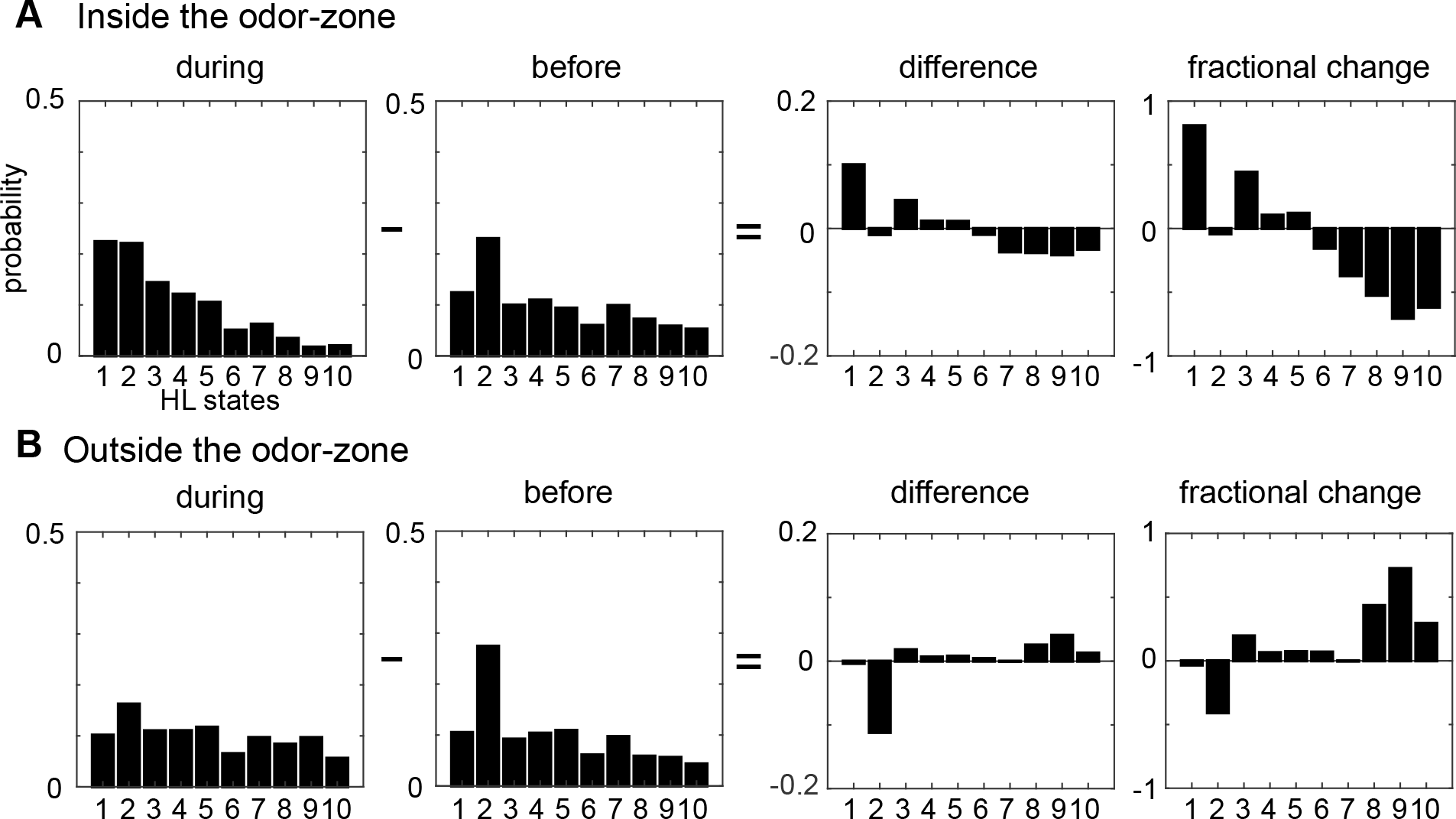
Odors modulate locomotion by altering the time spent in different HL states. **A.**Odor Induced changes in the occupancy of HL states inside the odor-zone. Bar graphs showing the probability distribution of the 10 HL states during odor, before odor,the difference between the two and the fractional change in the distributions when the fly is inside. **B.**Same as in A, but for outside the odor-zone.

In the absence of ACV, as expected, the state occupancy inside and outside the odor-zone are quite similar: The fly spends 30% of its time in state 2 and roughly equal time in all other HL states. Introducing ACV changed the fly’s locomotor behavior both inside and outside the odor-zone, but with opposite effects on the HL state occupancy inside and outside the odor-zone. Inside the odor-zone, in the presence of ACV, the fly spends more time in states 1 and 3 at the expense of time spent in states 7-10 (Figure 5A). These changes from high-speed states to low-speed states suggest that in the presence of ACV the fly is performing a local search, presumably to find food. Outside the odor-zone (Figure 5B), the fly spends more time in the high-speed states (HL states 8-10), with a decrease in the occupancy of HL state 2 (which includes stopping). Decreased stopping and increased high-speed walking with turning is likely to represent a different search strategy, wherein the fly might be attempting to re-find the odor it has recently lost. We also investigated whether there were changes in the LL state composition of the HL states and found that there were no changes (data not shown). Overall, these results show support for the second alternative above – odors, in our arena, affect locomotion not by creating new locomotor features, but by altering the frequency with which existing locomotor features are used.

The divergent effect of ACV on the probability of HL states inside and outside the odor-zone is consistent with our previous analysis^20^ and shows that the effect of ACV can be productively described by the change in the probability of the fly occupying HL states. To assess whether there is a more fine-grained spatial structure to the effect of ACV on a fly’s behavior, we divided the arena into a 60-by-60 grid and measured the ACV-induced change in the probability of occupying a given HL state at each of the 3,600 locations (Figure 6). Figure 6A shows the analysis for HL state 1. The probability that a fly is in state 1 increases dramatically only at the edge of the odor-zone (Figure 6A), and not throughout the odor-zone where the odor concentration is uniform throughout, showing that the effect of odor on locomotion has a fine-grained spatial structure. The color map representing a location-specific change in the probability of each HL state is shown in Figure 6B. The fine-grained modulation of locomotion is observed in other states as well - increases in state 2 are largest in an annular region just inside the odor-zone and increases in state 3 are largest at the very center of the arena. A similar specificity is observed in the increase in the probability of HL states outside the odor-zone. Increases in the occupancy of state 8 are uniform across the entire chamber outside the odor-zone; in contrast, the occupancy of states 9 and 10 increase in the region close to the odor-zone.

**Figure 6.**
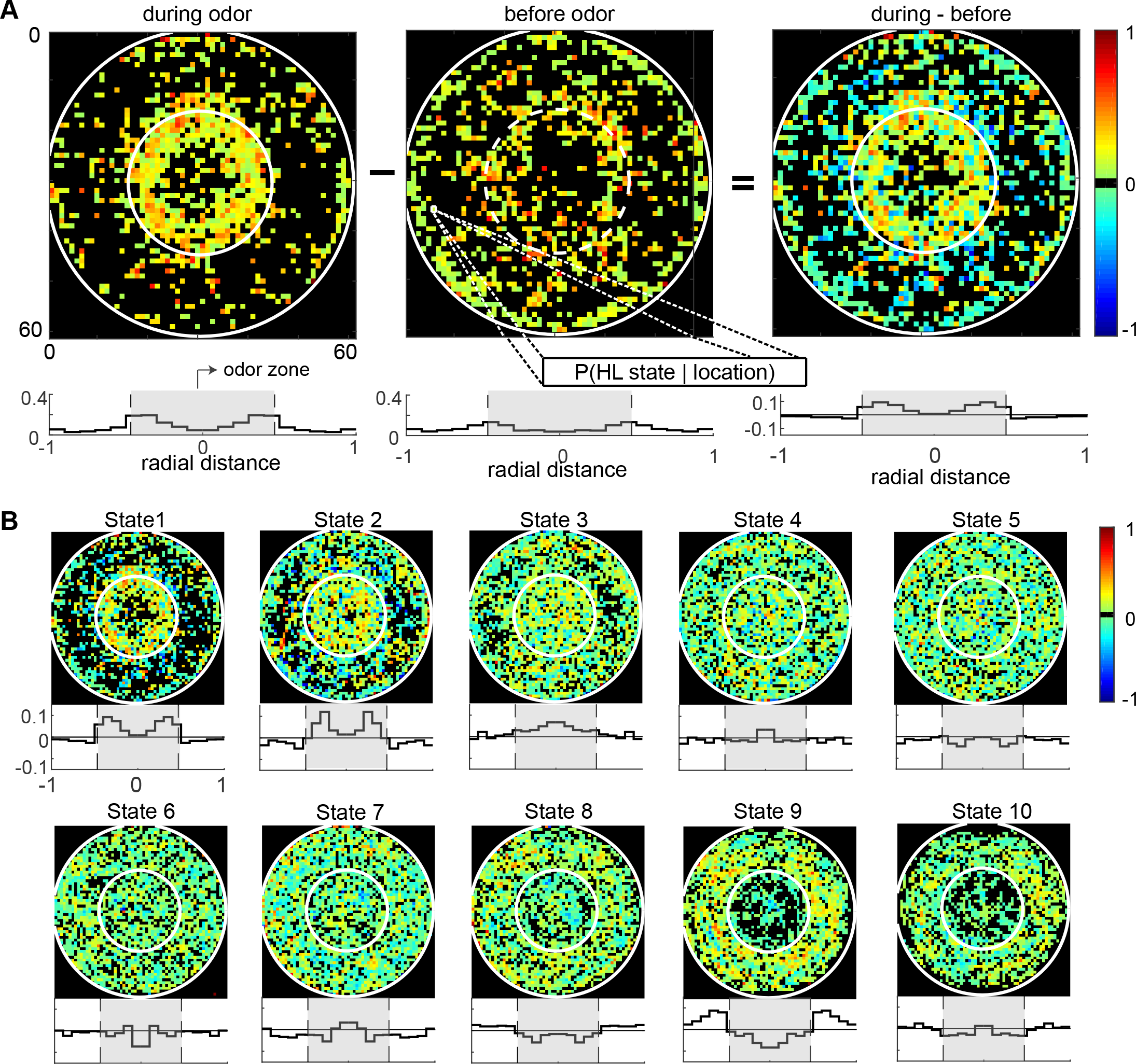
Fine spatial structure underlying odor-evoked changes in HL state occupancy. **A.**The probability distribution of HL states before and during odor was calculated for each of the 60×60 bins. Taking HL state 1 as an example, each specific bin represents the probability of HL state 1 given the spatial location. White circles show the extent of the odor ring and arena. Bottom row: Probability of HL state 1 as a function of radial distance away from the center of the arena is plotted for each scenario (bottom row). The shaded region indicates the bounds of the odor zone. **B.**Change in HL state occupancy for each of the 10 HL states are shown for based on 60×60 bins (top rows). Total change in HL state occupancy for each HL state as a function of radial distance away from the center of the arena (bottom rows).

In sum, the effect of odors on locomotion can be described accurately and concisely in terms of the change in the occupancy of the different HL states. One surprising insight from this description, unlikely to be discovered otherwise, is the fine spatial structure (Figure 6) with which odors modulate locomotion. One clear spatial structure that the flies employ when entering the odor-zone is to exhibit a stop and go behavior (state 1) when it enters the odor-zone because there is an odor gradient right at the border. Its stop-and-go behavior might relate to assessing the direction of increasing odor intensity. Once it is within the odor-zone its behavior is dominated by either a search behavior (state 3) or with relatively long stops (state 2).

### Both locomotion and an odor’s effect on locomotion is fly dependent

Given that ACV affects the occupancy of HL states, it should be possible to do the reverse, i.e., decode the presence of ACV based on the distribution of HL states. Surprisingly, a range of different decoding techniques failed to decode the presence of ACV based on the HL state. One such method (Figure 7-S1) in which we employed logistic regression to classify each 1 second of every fly’s track into “ACV present” or “ACV absent” failed. Even more surprisingly, population decoding based on HL states did not perform any better than decoding based on the observables (Figure 7-S1). One possibility that the logistic regression approach failed is because the average behavior represented in Figure 5 does not accurately encapsulate the behavior of individual flies. Large fly-to-fly differences, where different individuals are fundamentally different in their basal locomotion or their response to odor, might doom decoding methods aimed at discovering a single set of regressors that captures the behavior of individual flies.

**Figure 7.**
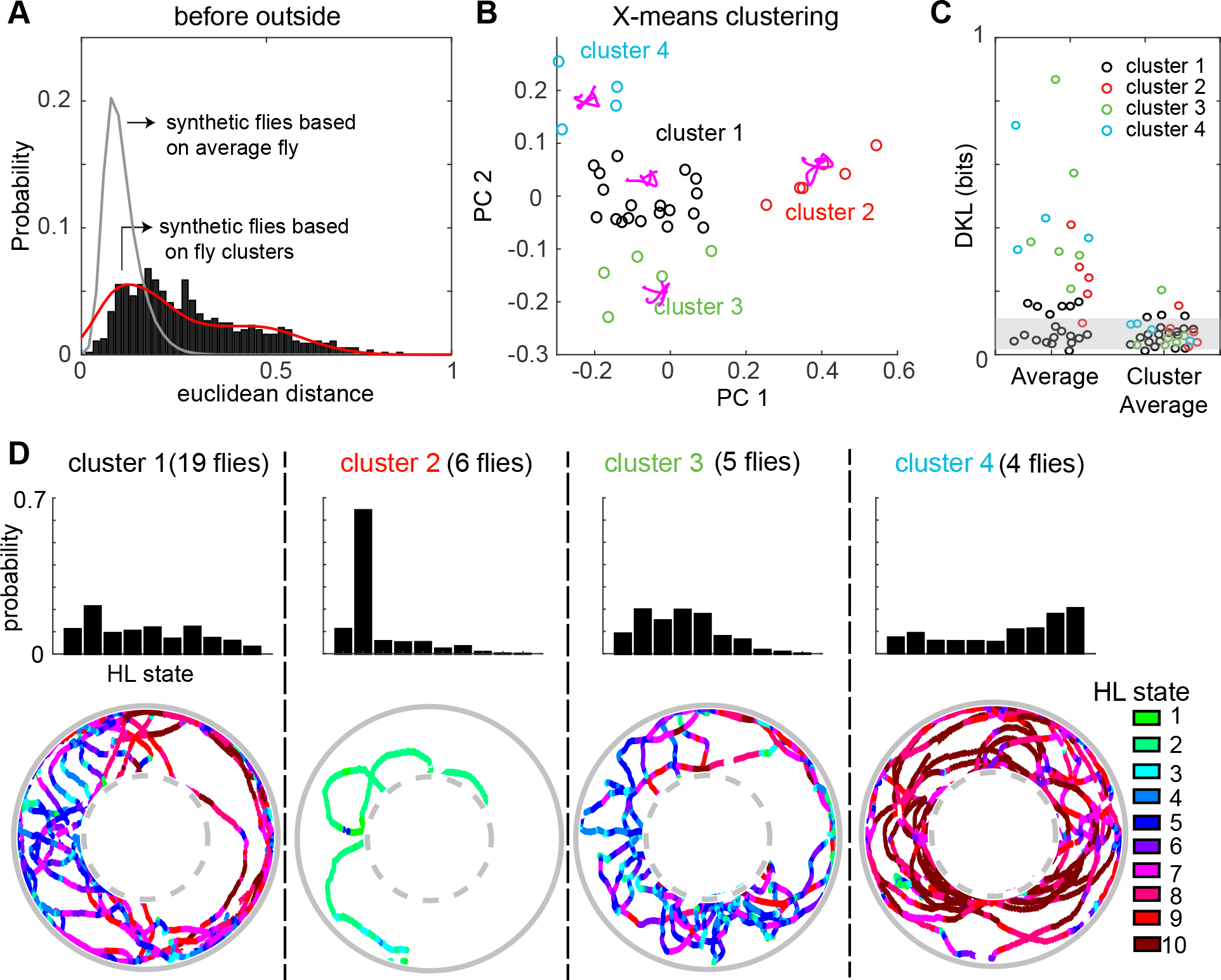
A fly’s locomotion cannot be explained as variations around an average fly and is well-described on the basis of 3-4 locomotor types. **A.**The empirical distances between flies in the 10-dimensional space formed by the HL states. The distance between 100 iterations of 34 synthetic flies based on the average distribution of states is much smaller (gray line, Wilcoxon rank sum test, p<10-131). The distance between synthetic flies drawn from 4 clusters of flies (red line) has a distribution more similar to the empirical distribution. **B.**X-means clustering (a variant of K-means) based on the 10-dimensional space formed by the HL states show that there are 4 clusters of flies in the “before odor/outside odor-zone” segment of the data. The first two PCs are shown for visualization. Each cluster is represented by a different color. **C.**Left: KL divergence from the HL state distribution of the average fly to the HL state distribution for each individual fly (n = 34). Right: KL divergence from the HL state distribution of the average fly in each X-means cluster to the HL state distribution for each individual fly in the corresponding clusters. Shaded area reflects the region within which 99% of KL divergences from the HL state distribution of the average fly to the HL distribution of synthetic flies generated from the average distribution (n = 3400). **D.**Average HL state distribution for each cluster is shown in the top row. Bottom row shows an example fly for each cluster with HL states overlayed. Colorbar for the 10 HL states is shown. A. An example of HL state as a function of time for a single fly. The HL states were segmented into 1s chunks and further subdivided into each of the 4 scenario - before inside, during inside, before outside and during outside. Here we only schematize Before Inside and During Inside because we show the logistic regression for the inside case. The HL state distributions were calculated for each segment. Thus each 1s bin is represented as a point in 10 dimensional state. We performed PCA on the entire dataset and the principal components that explains most of the variance (>90% variance) were used to fit to a logistic regression (logit) model. B. (Left) Results of a logrithmic regression (logit) model fit to the HL state distributions in 1 second segments for all flies do not show better probability of correct predictions over change (gray line). (Right) Distributions of high level states of the population do not improve decoding over observables.(Wilcoxon signed rank test,p<0.0084 for inside and p<0.0984 for outside).

As a first step to evaluating the extent of fly-to-fly differences in locomotor behavior, we measured the Euclidean distances between the representations of flies in the 10-dimensional space created by the 10 HL states. The distribution of distances between empirical flies was compared to the distance distribution between synthetic flies that are each drawn from the average fly (see Figure 7-S2 and methods section 8 for details regarding generating synthetic flies). We found that the distances between the empirical flies are much larger than the distances between the synthetic flies (Figure 7A). It is statistically impossible (p<10^-131^) that the observed Euclidean distance represents variations around the same average fly. Figure 7A shows the analysis for the before odor/outside odor-zone condition. The same conclusion applied to the fly’s behavior in the other three conditions – before-inside, during-outside and during-inside (Figure 7-S3).

Because the data in Figure 7A is inconsistent with variations around a single average fly, we assessed whether the observed variability – estimated by the Euclidean distances histogram in Figure 7A – can be approximated based on a small number of discrete locomotor-types. In Figures 8 and 9 we will assess whether dividing flies into a small number of discrete clusters will help decoding methods aimed at dividing the tracks into “odor” or “no odor” conditions.

**Figure 8.**
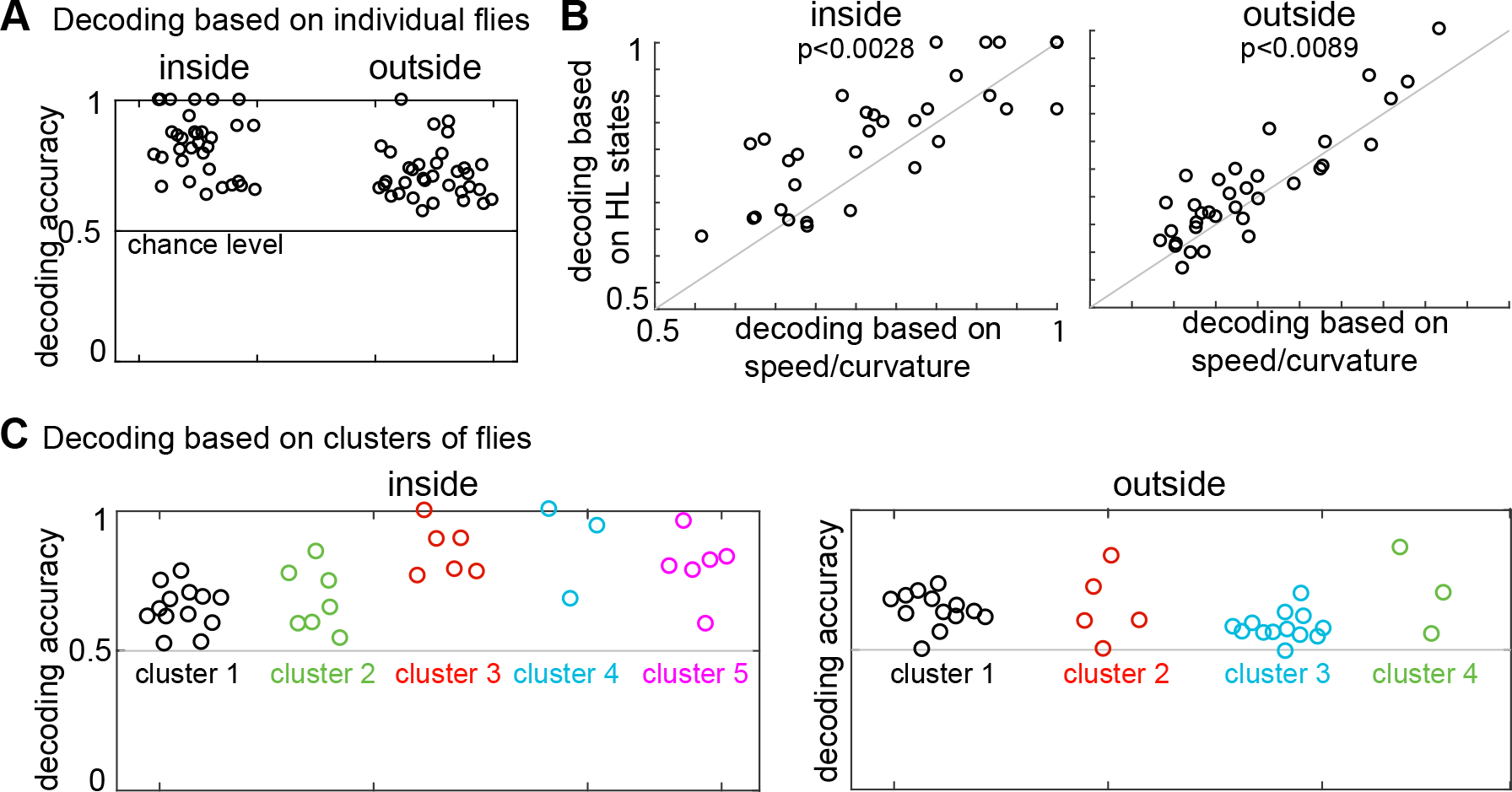
Presence of odor can be decoded based on both HL state distribution of individual flies and clusters of flies. **A.**HL states are a good predictor for experiencing odor for individuals. Results of a logrithmic regression (logit) model fit to individual flies consistently shows better probability of correct predictions over chance (grey line). **B**. HL states have higher predictive power than speed/curvature for individuals. Logit fits based on HL states were compared to that fit to the speed and curvature for inside (left) and outside (right) of the odor ring. A two-sided Wilcoxon signed rank test showed improved decoding using the HL states (p<0.0028 for inside and p<0.0089 for outside). **C.**X-means clustering (a variant of K-means) based on the 10 dimensional space formed by the HL states show that there are 5 clusters of flies for difference in the HL state distributions inside and 4 clusters of flies for difference in the HL state distributions outside. Results of the logistic regression model fit to these clusters for the inside and the outside cases are shown. Each cluster is represented by a different color.

**Figure 9.**
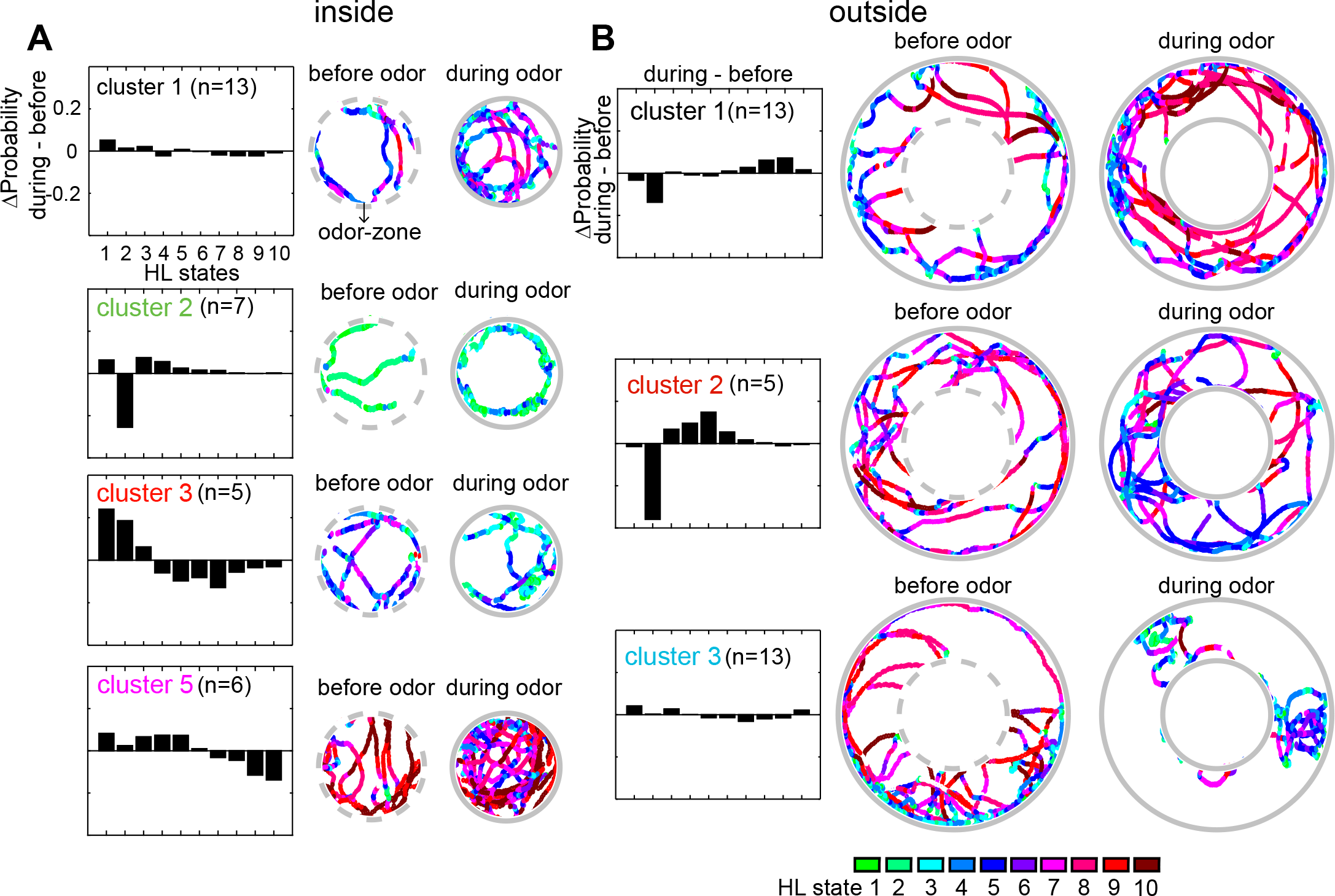
Flies’ response to odors cluster into a small number of distinct, sometimes opposing response-types. **A.**The 4 clusters with more than 3 flies differ in the change in states mediated by ACV. The difference between HL state distributions (during-before). Tracks from an exemplar fly from each cluster is shown. Dotted grey circle indicates the odor-zone. **B.**Same as A, but for odor-evoked changes outside the odor-zone. The colorbar for the 10 HL states is shown.

To test whether there are discrete clusters of lcoomotor-types, we performed X-means clustering (see methods section 6) and found that there are 4 clusters of flies based on their locomotion outside the odor-zone, before odor onset (Figure 7B). Although the identity of flies that cluster together changed, a similar number of clusters was found in each of the 4 conditions (Before odor/inside odor-zone, during odor/inside odor-zone, before-outside and during-outside, Figure 7 and Figure 7-S3). Importantly, X-means clustering on a set of 34 randomly sampled points from a uniform distribution in the probability simplex space that the data reside in found no clusters. If individual flies belong to 3-4 different clusters, we should be able to replicate the empirical distribution (Figure 7A) by creating synthetic flies modeled on 3-4 average flies, one representing each of the 3-4 different clusters. Indeed, the Euclidean distances between synthetic flies drawn from the 4 different clusters were similar to the distances between empirical flies (Figure 7B and Figure 7-S3). Since X-means clustering tend to underestimate the actual number of clusters in data^32^, the analyses in Figure 7 and Figure 7-S3 suggest that there are at least 3-4 fly-types based on the frequency with which they use the HL states.

Another method to assess the inaccuracy of representing individual flies with the average fly is to use an information-theoretic approach by computing the Kulback-Liebler divergence (KL divergence) to measure the amount of information loss when the average fly distribution of HL state is used to approximate the individual fly distributions of HL states. We measured the KL divergence from the average HL state distribution for all flies to the HL state distribution for each of the flies (Figure 7C). Consistent with the analysis with Euclidean distances above, the KL divergences (Figure 7C, left set of data points) show a large range indicating that some flies are well-represented by the population average while others are not. Does employing 3-4 fly-types defined as average distribution of their respective cluster decrease the information loss? Indeed, as expected we found that the information loss when we assume fly-types is much lower (Figure 7C, right). To place these KL divergences in an appropriate context, we also evaluated the KL divergence from the average fly to the synthetic flies generated from this average. This analysis shows that invoking 4 fly-types and describing the data based on these 4 fly-types is a demonstrably more accurate than describing it based on a single average fly-type.

How is the behavior of the flies in the 4 different clusters different from each other? The average occupancy of the HL states for flies in each cluster and one example from each cluster is shown in Figure 7D. One cluster of flies (cluster 2) is distinct from the others because flies in this cluster move at markedly slower speed than the rest; flies in cluster 2 spend >60% of their time in State 2 (Figure 7D) during which the fly is often stopped. Locomotion of flies in the largest cluster (cluster 1) is characterized by an alternation between medium-speed states and slow-speed states. Flies in this cluster employ states 5 and 7 with a high frequency while making radially-inward forays into the center of the arena. Similar behavior is observed in another cluster (cluster 3), except that the flies employ the slower medium-speed states (states 4 and 5). Finally, the fourth cluster of flies demonstrates a different locomotor strategy. Instead of radial forays to the center of the arena that characterizes the behavior of flies in clusters 1 and 3, flies in cluster 4 traverse the arena in concentric circles using the high-speed states (states 8-10). Thus, X-means performed on the HL state distributions appear to identify different locomotion strategies employed by the fly.

That the observed distribution of Euclidean distances between flies is better explained by clustering the flies into 3-4 locomotion types than by assuming that there is a single average fly suggests fundamental fly-to-fly differences in locomotion, and partly explains why decoding based on the average state distribution fails (Figure 7-S1). Another reason is that the behavior of the fly before odor onset is only weakly predictive of its behavior during the presence of odor.

First, consider flies such as flies 11 and 33 whose behavior in the before odor/outside odor-zone is similar to each other (Figure 7-S4A, see the distributions of HL states), their locomotion during the presence of odor is different enough such that they are in different clusters (Figure 7-S4A). Flies 17 and 27 represent another pair whose behavior before odor onset is similar (Figure 7-S4A), but behavior during odor is markedly different. A similar trend is observed inside the odor-zone (Figure 7-S4B). These examples imply that behavior before odor onset is not predictive of behavior during the presence of odor. To evaluate the similarity between a fly’s locomotion before and during odor, we measured the correlation between Euclidean distances between flies in the before and during periods. If the behavior of the flies before odor-onset is predictive of the behavior of the flies during the odor, we should observe a strong correlation between the two. Instead, we find that the correlation is rather weak (Figure 7-S4A and Figure 7-S4B).

### A small number of strategies can explain the variability in flies’ response to odor

The analysis presented above suggests that individual differences explain why the logistic regression approach based on the average HL state distribution across flies failed to decode the presence of ACV from its absence based on the HL states usage by individual flies. If so, individualized logistic regression aimed at decoding whether a given fly is experiencing the presence and absence of ACV at any time based on the distribution of its own HL states should be more successful. Indeed, logistic regression based on individual flies: where regression was performed to decode the presence or absence of odor based on HL state occupancy during a 1s-interval was able to perform an odor-no odor classification at better than the chance level for every single fly (Figure 8A, see methods section 7 and Figure 8-S1 for details). Moreover, as expected, logistic regression using the HL states performed significantly better than did the observables (Figure 8B) implying that HL states are more predictive of the presence of ACV than the observables.

The analysis in Figure 8A shows that the occupancy of HL states is predictive of the presence or absence of odor when the analysis is performed at the level of individual flies. Does this mean that each fly follows an individualistic strategy? Given that there are 3-4 clusters of flies (Figure 8 and Figure 8-S3) both before and during the presence of ACV, we anticipated a similar number of clusters or strategies underlying the change in locomotion due to ACV. To evaluate whether a fly’s response to ACV cluster into a small number of response types, we once again started with X-means clustering based on the change in state occupancy before and during the odor. Two separate X-means clustering were performed – one for odor-evoked differences inside the odor-zone and another for odor-evoked differences outside the odor-zone. X-means clustering found 5 clusters inside the odor-zone and 4 clusters outside the odor-zone (Figure 8C). Using these clusters as a starting point, we could reconfigure the clusters such that the logistic regression on flies in each cluster performed at a better than chance level for each fly in the cluster (Figure 8C, see methods section 7 for details), thus implying that a fly’s response to odors can be approximated as a choice between a small number of response-types.

To understand the differences between strategies that flies employ to alter locomotion, for each cluster, we plotted the difference between mean state occupancy before and during odor exposure (Figure 9). There were four clusters with more than three flies for inside the odor-zone (Figure 9A). Flies in cluster 5, just like the average fly (in Figure 5), slow down inside the odor-zone. Flies in cluster 3 also demonstrate a strategy similar to flies in cluster 5 except that ACV causes a large decrease in the time a fly spends in the medium-speed states rather than the high-speed states. In contrast to clusters 3 and 5 during which the fly slows down inside the odor-zone, flies in cluster 2 demonstrate a fundamentally different strategy in which there is a large decrease in state 2 in favor of states 1, 3 and 4. The flies in this cluster go from stop-start locomotion to locomotion in which they either meander at slow speeds or walk slowly with many sharp turns. Finally, for the flies in cluster 1, there is no dramatic change in state. These different strategies represent diametrically different effects of ACV on some HL states – the most striking example is the opposing effects of the odor on HL state 2 occupancies in different clusters – a large decrease in cluster 2, and an increase in clusters 3 and 5. These differences explain the odor-induced increase in usage of all the slow states, except state 2, inside the odor-zone for the average fly (Figure 6A).

Outside the odor-zone, flies respond to ACV in three distinct ways (Figure 9B). Flies in cluster 1 decrease the time they spend in slow-states (state 1-2) and instead spend time in the fast states (states 8-10) resembling the behavior of the average fly. Another cluster of flies’ behavior (cluster 2) shows a large decrease in HL state 2 occupancy similar to the behavior of flies in inside odor-zone cluster 2 while exhibiting a large increase in medium-speed states (state 4-6). Finally, the behavior of the third cluster of flies (cluster 3) show no dramatic change in state. In sum, just like locomotion before the onset of ACV, ACV has different effects on locomotion in different flies. Both locomotion and the effect of odors on locomotion on individual flies are much better explained by dividing the flies into a small number of clusters than as variations around a single average fly.

## Discussion

A cornerstone of neuroethology is that behavior unfolds in discrete packets, i.e., behavior can be temporally segmented into natural units^33–35^. In some behaviors, these discrete packets are readily recognizable. But, in most daily behaviors, there is enough variability in these discrete packets to make the underlying natural units unrecognizable without the help of sophisticated analytical tools. In the last few years, there has been significant progress in discovering structure in sequences of posture in *C. elegans17*, in *Drosophila*^9,12^ and in mice^11^. In this study, we find a similar structure in the description of fly’s locomotion in the velocity space.

Our salient results are:

- A fly moves at a relatively constant 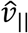 and 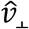 for tens of step. This tendency means that a fly’s locomotion can be decomposed into a small number of locomotor features – 10 features in the case of the model we present. The fact that the same 10 locomotor features can describe the behavior of all the flies in our dataset is unexpected, and allows a simple description of the effect of odors on locomotion, and of the differences in behavior across flies in terms of how these 10-locomotor features are employed.
- Using this analytical framework, we show that odors affect locomotion by altering the time that a fly spends performing a given locomotor feature (instead of creating new features). The odor-induced change in locomotion has a fine spatial scale – the fly’s response to odor changes as it moves from the border of the odor-zone to its center, and as it moves away from the odor border.
- The HHMM framework also allowed us to show that although different flies in our dataset used the same 10 locomotor features, they used it in different proportions. The variation is so large that the fly’s behavior cannot be understood as variations around the same average fly. Instead, the flies employ a minimum of at least 3-4 different strategies.

Below we discuss the limitation and implication of these findings.

### Limitations of the model

The model presented here is <u>*a*</u> model of locomotion and not <u>*the*</u> model of locomotion. The choice of observables and model will strongly influence the features of the structure that is discovered. Our particular model reveals the structure of locomotion in the velocity space.

In choosing the observables, we employ a common method for describing locomotion, i.e., we treat the fly as a point object and measure the instantaneous change in the position of this point object; therefore, much of the insights from the model relate to how the fly changes its position in time. Apart from 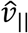 and 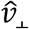, another similar and more commonly used representation of the change in fly’s position: instantaneous speed and angular speed yielded similar locomotor features (data not shown). Ultimately, we used 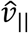 and 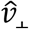 because this representation is more closely related to movement representation within the insect brain^36–38^, and because the measurement errors associated with angular speed are particularly large when the fly is moving slowly^24^.

A fly’s position in the world can also be described using the actual position of the animal as observables rather than the change in the position, as has been employed in a recent study in rats^39^ Using the instantaneous positions as observables will reveal different aspects of the structure underlying an animal’s locomotion: One example is the trajectories of flies in Clusters 1 and 3 Figure 7. They cross similar spatial positions but are classified into different states because the flies are traveling at different speeds. In terms of the sequence of position in space, flies in both clusters have a similar behavior – they explore outer arena border and make occasional radial forays inside the odor-zone. An analysis based on position would likely place these two clusters of flies together whereas our analysis, in the velocity space, places them in different clusters.

Model architecture is also important. A hierarchical model performed better than a non-hierarchical model. The current model has state durations of <3 seconds. It is clear to human observers that there is structure in the data that is > 3 seconds long. Flies sometime explore the outer border of the arena using characteristic paths that can last up to a minute. The short duration of states in our model cannot capture structure on these long-time scales. One possibility is choosing a deeper-layered architecture because given the structured transitions between the HL states in our model; it is likely that if we used a deeper-layered architecture, we would likely uncover structure on a longer timescale.

### Locomotor features and implications for neural control of behavior

One surprise in this study is the extent to which a fly’s locomotion is structured in the velocity, both in terms of the narrow distributions of 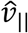 and 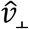 which describe each HL state, the persistence of each state and transitions to states that occupy neighboring regions in the velocity space. Consider the subtle differences between HL states 1 and 2. During both HL states, the fly’s locomotion is quite slow, but in state 2 the fly stops and runs intermittently while in state 1, the fly is continuously in motion, albeit slowly. Similarly, in each of the HL states 4, 5 and 7, 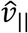 lies within a narrow range, which is distinct for each of these three states, implying a tight control over forward speed. These locomotor characteristics can persist over 3 seconds – a time period during which a fly takes 30 steps on average (given a step frequency of 10 Hz^40^). This tight control over 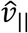 over tens of steps strongly suggests that locomotion unfolds, not on a step-by-step basis, but in blocks of tens of steps.

Another surprising result is that the same set of locomotor features describes the behavior of all the flies in the dataset. This result is particularly surprising given that our model explicitly allows each fly its own set of locomotor features (see methods). The fact that all flies can be reasonably modeled by the same model implies that within a given environment all flies construct their locomotion from the same building blocks, and differences in locomotion amongst flies or the effect of sensory stimulation can be quantified as changes in the frequency with which these building blocks are employed.

However, we anticipate that locomotor features are unlikely to be fixed. The particular set of HL states that we describe here represents the locomotor features that a fly uses as it explores a small, circular arena in the dark. The large proportion of left and right-leaning trajectories (States 8, 9 and 10) are likely a result of the circular shape of the arena; we are unlikely to observe these locomotor features in a long, rectangular arena. We contend that in any given environment, the fly’s locomotion is constructed from a small number of locomotor features that depend on the shape and size of the arena (among other things such as state of the fly).

Based on our results, we speculate that the statistical structure revealed by the HHMM is a reflection of the mechanism by which locomotion is controlled by the brain and suggests that the circuits in the brain must perform three tasks. First, given an arena and other sensory conditions, flies (and other animals) construct their search strategies from a small number of locomotor features. Second, based on an individual fly’s evaluation of its environment, it uses different locomotor features in different proportions. Finally, circuits that perform sensorimotor transformation, in large part, affect locomotion by altering how the animal chooses different locomotor features.

### Odors affect behavior on a fine spatial scale

In nature, animals encounter odors in a cluttered and dynamic sensory environment^41^. Discriminating between odors, navigating towards the chosen odor source and pinpointing the source of that odor requires a flexible deployment of multiple different motor programs. It is difficult to replicate the complex natural environment in the laboratory. Therefore, laboratory studies are aimed at different subsets of the complex environment experienced by animals. In insects, much of the work has focused on one such environment – an environment in which it experiences odors in a highly structured odor plume often within a high-contrast visual environment^42–45^. Recently similar experiments have been repeated in the context of flies walking towards an odor source^46^. These experiments seek to model an insect’s behavior in one specific condition in which it tries to locate an odor source at a distance using strong directional information from wind and vision. The experiments described here explore a fly’s behavior in a small circular arena. Moreover, we have eliminated access to visual information, and have minimal, likely undetectable wind cues^20^. Thus, this study models a fly’s behavior near the source of odor under conditions in which odor cues are the salient sensory cue.

We find that this behavior near the odor source can be productively described by changes in the HL states. The clearest evidence that changes in HL states are a good description of the fly’s behavior is the analysis in which we measured the spatial distribution of odor-evoked changes in HL states Figure 6, and observed a pattern that strongly resembles the odor-zone. One surprising insight from this description is that odor modulation of locomotion has a spatial scale that is smaller than the odor-zone. In the absence of odor, as expected, an average fly has a similar distribution of states both inside and outside the odor-zone. This basal behavior is spatially structured by the presence of ACV both inside the odor-zone and outside it. As the fly enters the odor-zone, it immediately transitions to state 1 during which the fly moves slowly without long stops. As it enters further inside the odor-zone, it transitions to state 2 during which short, low-speed movements are punctuated with complete stops. At the very center of the arena, the fly transitions to state 3 during which the fly’s locomotion is described by slow, sharp turns. The distinct radial distribution of the 3 slow states is unexpected and implies that the fly has a specific search strategy as it explores the odor-zone.

A similar spatial structure in the modulation of states is observed outside the odor-zone, although this structure is not quite as clear. There is a large increase in the occupancy of state 9, and to a lesser extent in state 10, immediately outside the odor-zone. The increase in the occupancy of these states during which the fly is walking at its fastest imply that these states are recruited as the fly exits the odor-zone, and results in a decrease in the time it takes for the fly to return to the odor-zone. We expected symmetrical change between state 9 and 10 but found that the change in state 9 is greater than the change in state 10. This difference is not due to the handedness of different flies^47^; rather it appears that most flies that use state 10 also use state 9 but to a lesser extent. In contrast to the increase in state 9 and 10, the increase in state 8 occurs throughout the area outside the odor-zone and might reflect an overall increase in a fly’s speed in the absence of odor.

### Flies show considerable variability in locomotion despite employing the same locomotor features

There is a growing recognition that even single-cell organisms and animals with simple nervous systems display substantial individuality^48,49^. Animals with larger nervous systems are likely to display even greater individuality, and in the case of the adult flies this individuality has already been demonstrated in the context of locomotor handedness^47^ in a choice assay. The nature and extent of individuality is harder to assess in more complex behaviors because of the difficulty in assessing individuality in a large behavioral space; differences in behavior can be either normal variation or fundamental differences in behavior. In this study, we find that different flies employ the same locomotor features but use them in vastly different proportions. Based on the distribution of HL states, we were able to demonstrate unequivocally that the observed variability between the flies is inconsistent with a single type of locomotor behavior but can be approximated by invoking 3-4 clusters of flies. Because the clustering framework we employed (X-means) underestimates the number of clusters, and because there were only 34 flies in our dataset, it remains to be seen whether there are indeed a small number of locomotor-types or a whole continuum of locomotor-types. Another limitation of the current study is that we have not yet ascertained whether the behavior of an individual fly persists over time. Despite these limitations, this study makes an important contribution to the study of individuality in two ways. The first contribution is by developing a statistical framework for the study of complex behaviors. This method can be extended to understand whether the differences we observe are true individuality. Second, in most behavioral studies, researchers are focused on understanding the effect of some stimuli on behavior. In these studies, conclusions are invariably based on the average fly. In this study, we provide a framework for testing whether the description based on an average fly is appropriate, and ways to proceed if such an approach is inadequate.

What is the general strategy that characterizes each of these clusters as the fly explores the arena before odor onset? Before odor onset and outside the odor-zone, 55% of the flies are in a single cluster Figure 7 – flies in this cluster either perform stop-start locomotion (state 2) or are moving straight ahead at a medium speed (state 5 and 7). Flies in another cluster display similar behavior, except that they choose a slower speed (using states 4 and 5). These flies spend much of their time at the outer arena boundary and make occasional forays to the center of the arena. The flies in the other two clusters have fundamentally different strategies. Flies in cluster 2 spend a much larger proportion of their time stopped, often away from the arena border. Finally, the fourth cluster of flies circles the arena instead of making radial forays to the center of the arena and thus make greater use of states 9 and 10. Thus, our analytical framework identifies patterns which are quite apparent to the observer after the fact!

Flies’ behavior in the presence of odor-zone is similarly diverse. A simple zeroth-order description of the average fly’s behavioral response to ACV, without the nuance we discuss here, would state that flies respond to odors by employing the lower speed states to a greater extent. Even this zeroth level description only fits half the flies in our dataset (Clusters 3 and 5, Figure 9A). Many of the rest of the flies respond by increasing their speed rather than decreasing it. These differences are not correlated with attraction index^20^, a measure of a fly’s attraction to an odor. Thus, flies can flexibly deploy different locomotor features to achieve the same attraction index. The behavior of different flies outside the odor-zone is similarly diverse. For some flies, the fly’s behavior inside and outside the odor-zone is remarkably similar and consistent with a slow, local search implying that they disregard the odor boundary. In contrast, the behavior of other flies is highly sensitive to the odor boundary, and their behavior inside and outside the odor-zone is diametrically opposite: Inside they walk slowly, stopping frequently and turning sharply, presumably to find the source of the odor. Outside they walk fast in smooth trajectories that lead them back inside the odor-zone. Finally, there are flies that cannot be described by either of these two behaviors but by a combination.

The diversity of odor responses observed here is consistent with work done on moths where similar diversity of behavioral responses was observed^50^ but in sharp contrast to recent work on walking Drosophila^46^ in which the authors reported that attraction to odors results from a stereotypical motor pattern. The authors of that study claimed that the relatively simple response to odors in their study is likely a result of their simple behavioral arena. Another possibility is that in their study there is a strong, directional wind cue. In the presence of a steady wind cue in a narrow arena, it is likely that the flies’ locomotor behavior is dominated by upwind walking and suppresses other elements of their behavior. It is well-established that a fly’s response to odor is strongly influenced by context, as was demonstrated recently by comparing the response to odor in different visual and air flow conditions^51^.

Until recently variability has been shunned by neuroscientists and often treated as noise in measurements. A series of elegant studies in the context of stomatogastric ganglion in lobster has established that properties of neurons that are part of a circuit which produces a simple rhythm are quite variable^52–54^ leading to an appreciation of variability in neurons followed by the similar discovery in other systems^55^. A similar rethink is underway in the context of behavior. Any scientist who has attempted careful behavioral experiments knows the considerable variability in behavior. When considered from the viewpoint of an individual animal, this variability is hard to understand: A hungry fly in search of food should respond with a singular, hardwired behavior which represents an optimal strategy for locating food. However, species evolve as large populations of individuals, and a successful species should be able to adapt to fluctuating environmental condition. Recent work has shown that - bet hedging - a process by which the same genotype shows considerable phenotypic variation is an important strategy for adaption to fluctuating environment^56^. The basic idea is that having a diversity of phenotypes ensures that some individuals would thrive in any condition. Thus, variability might make sense in the context of species. We think that behavioral variability is a feature not a bug and its careful consideration is critical.

The study of an individual’s behavioral response is also important to understand the mechanism underlying both the control of locomotion and how odors control locomotion. Analyzing behavior at the level of population can provide important insights into an animal’s response but does not provide the resolution necessary for understanding the neural mechanism underlying the moment-by-moment control of behavior at the level of individual flies. In this context, it is instructive to take a closer look at the average response to odors in the light of clusters of response to odors. The average response Figure 5A has an odd feature–the occupancy of all the slow states except state 2 is increased. The lack of increase in occupancy of state 2 results from a cluster of fly in which the occupancy of state 2 is strongly decreased. Figure 9A. A similar effect is observed in the response of the average fly to the medium-speed states – states 4-7. The occupancy of these states decreases in some flies and increases in other flies. Thus, the average fly is an aggregate of these different clusters of flies, each of which has a distinct response to odor. Disaggregation is an essential first step to understanding neural control of behavior.

## Materials and Methods

### 1. Collection of behavioral data

The methods used to collect the behavioral data were reported in a previous study^20^. Briefly, flies were raised in a sparse culture. Flies that were 3-5 days posteclosion were starved for 14-18 hours. Locomotion of a single fly was recorded for a 3 minute period before odor was introduced (before period) and a 3 minute period during odor (during period), using a video camera at a rate of 30 frames per second. The coordinates of the fly were extracted using a custom Matlab program.

### 2. Extracting observables from the trajectory

The behavioral arena was normalized to a unit circle centered at the origin. The raw coordinates of the centroid of the fly were smoothed using a wavelet denoising algorithm followed by a locally weighted (lowess) algorithm. Speed and curvature was defined exactly as in the previous study.

To quantify the behavior of the 34 flies, we computed the speed of the fly along the original direction of movement 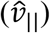 and the speed of the fly perpendicular to the original direction of movement 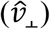. 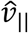 at time *t* was defined as the component of the velocity at time *t* in the direction of velocity of the fly at time *t* − 1. 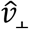 was defined as the component of the velocity at time *t* perpendicular to velocity of the fly at time *t* − 1 (Figure 1C). These values were calculated as follows:

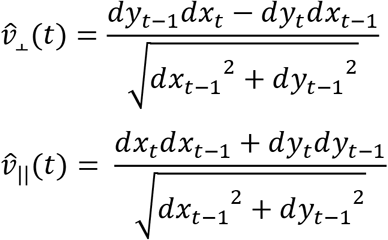

Values of 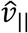 and 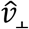 found to be further than 4 standard deviations away from the average were set to values drawn from a normally distributed distribution (*σ* = 1) centered at the 4 standard deviation mark. The parallel and perpendicular velocities were then set as the observables used in fitting a 2-level Hidden Hierarchal Markov model (HHMM).

We employed 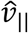 and 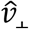 instead of speed and curvature because curvature is very noisy at low speeds because the calculation of curvature requires division by the third-power of speed. These two variables are directly related to speed and curvature:

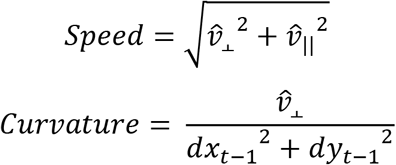

### 3. HHMM modeling

HMMs are widely used in a variety of fields for modeling time series data. An HMM is a markov model which assumes that a given sequence of observations may be explained by a set of non-observable states and the time independent probability of transitioning between these states. The model processes which produces the observations in an HMM are hidden to the researcher and thus, the goal of fitting an HMM is to uncover the highest likelihood probability model parameters that can generate the data. Baum and others developed the core theory of HMMs^57^. Since then there has been much exploration of model architecture, and fitting procedure. The HHMM is an extension to the HMM which applies hierarchical structure in the form that higher level state is in of itself an HHMM composed of its lower level states.^23^ The approach we take of exploring and fitting the HHMM closely follows the approach developed by Matt Beal in which he applied variational algorithms to fit HHMM to a time series of observables^58^.

This section is divided into three parts. First is the description of the model, second is the details of the process by which the model is fit, and third is the thought process behind our model selection.

#### 3.1. Model description

The model we describe here has 10 hidden states at the higher level (HL) and 5 hidden states at the lower level (LL) Figure 2A. The empirical data to which the model is fit is a time series of the two observables - 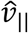 and 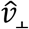. In building the model, the aim is to assign each instant in this time series to a LL and HL state. Each LL state is associated with a joint probability distribution on the observables. Each HL state is described by the transition probability matrix of its LL states. Therefore, while fitting a HHMM, we are determining three sets of quantities: First, the distribution of 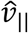 and 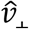 which describes each LL states (Figure 3-S1A/B). Second, the transition probabilities (TP) between the LL states which describe each HL state (Figure 2-S2B). Finally, the transition probabilities between the HL states (Figure 2-S2A). Based on the values of 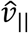 and 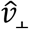, the model assigns a sequence of HL and LL states which best describes the data.

TP matrix for the HL states and the LL states associated with each HL states is shown in Figure 2-S2. In each case, the TP matrix describes the probability (Pij) that a fly in a given state detailed in the *j*^th^ column will transition to a different high-level state detailed in the *i*^th^ row. The high-level states are arranged in ascending order of mean speed/variance in curvature ratio. Because of this arrangement, the TP for the HL states appears well-structured. From any state, there is a strong tendency to transition to one of the neighboring state. This tendency implies that flies transition to states which have a similar speed/curvature ratio. LL states that belong to the same HL state often have more similar speed/curvature ratios. Thus, there is less of a tendency for LL states to transition to the LL state with the closest speed/curvature ratio. Nevertheless, the LL state transition probability matrices are sparse; signifying a distinct pattern of transition between LL states (Figure 2-S2B).

Figure 3-S1A/B shows the joint distribution for 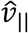 - 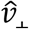 for each state. Each row represents one HL state and its low-level children. In each panel, the solid line represents the model predictions and dots represent empirical values. Each LL state is modeled as a multi-variate normal distribution of observables. The solid line represents the bound within which the model predicts that 85% of the data points corresponding to a given LL state lie. The empirical data points represent randomly selected subset of data corresponding to instants which the model assigns to a given LL state. The close agreement between the two for most LL states implies that the model is an excellent descriptor of the observables. The few exceptions where either the model distribution is too broad or where the model is not a good descriptor of the data reflect cases in which there are not too many data points in the concerned state (See for example, LL state 1 for HL state 6, Figure 3-S1A). Each HL state is a composite of the 5 LL state because for every time instance the fly is in a given HL state, the model assigns a LL state. Therefore, we can consider the probability density function of a HL state as the sum of the probability density functions of its LL states. From this, we can calculate the 85^th^ percentile contour for the corresponding probability density function for the HL states (Figure 3A2, Figure 4). Again there is a strong agreement between model and empirical data.

#### 3.2. Fitting process

We will first formally define a HMM as follows:

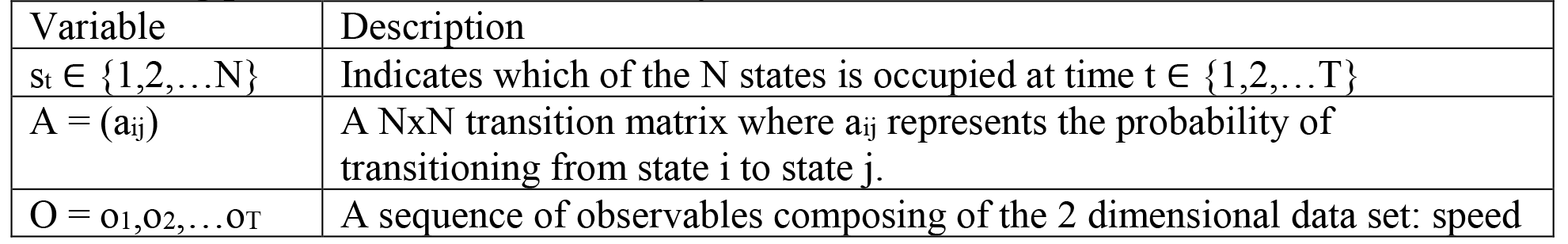

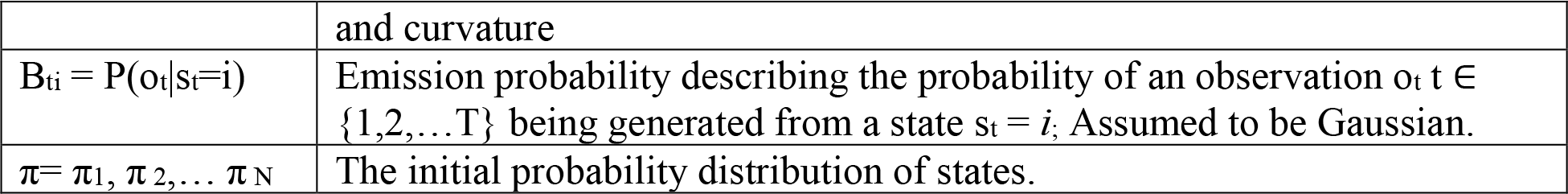

For our 2 layered HHMM, we chose a structure composing of 10 high-level states (HL state) on the top level with each HL state being associated with 5 low-level states (LL states) on the bottom level (Fig. 2). Each LL state is modeled as a multivariate normal (MVN) distribution on the observables, and the HL states as being fully described by the mixture of LL state distributions associated with the HLS.

Now we will define our 2 layered HHMM as follows:

| Variable | Description |
| --- | --- |
| $s_t \in \{1, 2, \dots, N_{HL}\}$ | Indicates which high level state is occupied at time $t \in \{1, 2, \dots, T\}$ |
| $u_t \in \{1, 2, \dots, N_{LL}\}$ | Indicates which low level state is occupied |
| $Q = \{q_{kl}\}$ | High level state transition probability matrix, $q_{kl}$ represents the probability of transitioning from high level state $k$ to high level state $l$. |
| $A^k = (a_{ij}^k)$ | The low level state transition probability matrix. Here $a_{ij}^k$ represents the probability of transitioning from low level state $i$ to low level state $j$ given that the high level state is $k$ |
| $O = o_1, o_2, \dots, o_T$ | A sequence of observables composing of the 2 dimensional data set: speed and curvature |
| $B_{tki} = P(o_t \vert s_t = k, u_t = i)$ | Emission probability describing the probability of an observation $o_t$ $t \in \{1, 2, \dots, T\}$ being generated from a HLS $s_t = k$ and associated low level state $u_t = i$. Assumed to be Gaussian. |
| $\pi^0$ | The initial probability distribution of HLS. |
| $\pi^k$ | The initial probability distribution of LLS for HLS $k$. |

We can henceforth refer to the set of model parameters as *θ* = {*π*, *A*, *B*} and the latent state variables as *Z* = {*s*_1_,*u*1, *s*2,*u*2, *s*3,*u*3 … *s*_*T*_, *u*_*T*_}. For a given set of observations the goal is to obtain a posterior probability distribution over the parameters and latent state variables. We approximate this posterior distribution as a factorized distribution over latent variables and parameters, i.e. *p*(*θ*, *Z*) ≈ *q*(*θ*)*q*(*Z*), and use the Variational Bayesian Expectation Maximization (VBEM) algorithm to find the best approximation. *q*(*Z*) is obtained using the forward-backward algorithm which provides sufficient statistics needed to update the approximate posterior distributions over the parameters. In this setting, posterior probability distributions over rows of transition probability matrices are assumed to be Dirichlet, and the prior is chosen to favor self-transition parameters, *α*_*ii*_ > 1, while discouraging the use of unneeded states i.e. *α*_*ij*_ < 1, for, *i* ≠ *j*. Initial state distributions were also assumed to be Dirichlet with *α*_*i*_ < 1. The emissions probability distributions associated with each state were assumed to be Normal inverse Wishart with a prior favoring zero mean and unit variance. For each computational run, the initial parameters of these posterior distributions were randomized.

##### VBEM algorithm

The Variational Bayesian Expectation Maximization algorithm functions by iteratively updating the two components of our factorized approximation (Equation 1) to the true posterior over parameters and latent variables and exploits conditional conjugacy to identify explicit update rules for all distributions over parameters. The goal of the VBEM algorithm is described alternatively as minimizing model error as given by the Kullback-Leibler divergence between the approximate and true posterior distributions; or maximizing a lower bound on the marginal probability of the data given the model. It is perhaps easiest to see how the algorithm works by considering the KL(q,p) instantiation. Under our factorized approximation this objective function takes the form

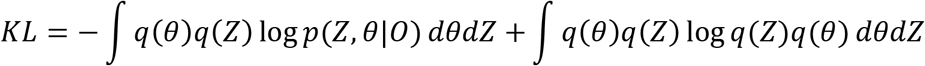

The VBEM algorithm minimizes this objective function using coordinate ascent in the q(θ) q(z) function space. That is we obtain iterative update rules by simply taking the functional derivative if the KL with respect to q(Z) for fixed q(θ) and then solving for q(Z). This results in the so-called E step where we update the posterior distribution over latent variables by averaging the true joint distribution of observations, parameters, and latent variables over our current estimate of the posterior on parameters:

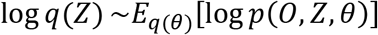

In the so called M step, the roles are reversed:

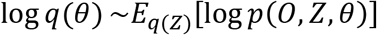

Here the tilde indicates equality up to an additive constant. This two-step procedure is repeated until convergence. The simplicity of these equations belies the complexity of the actual calculation of the posterior distribution over latent state variables. This is accomplished using the well-known forward-backward algorithm. The particular implementation of the forward-backward algorithm used in VB differs from the traditional EM implementation in that the parameters of the transition probability matrix are obtained by exponentiation of the geometric mean of the transition probability matrices:

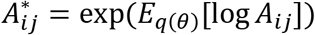

Based on the current posterior over parameters instead of simply using a maximum likelihood or MAP estimate. Because we assumed the rows of the transition probability matrix to be sparse, this further encourages the model to leave un-needed states unused.

Regardless, based on the assignments of observable tracks to HHMM models, we can iteratively update model parameters for each cluster to maximize the probability of observing the observables given the model parameters and the initial LLS probability density using the forward-backward algorithm. This operation can be thought of as the maximization step where we are calculating the best set of model parameters (*θ*) that maximizes the Q function.

Although the EM algorithm is guaranteed to get a better fit on every iteration, it often does not converge to a global optimum of the likelihood function. As such, we implemented a system of cluster pruning, splitting, and reassignment to perturb the system in the case of reaching a local minimal fit. In the pruning step, unused clusters are removed until we are left with at most 2 unused clusters. In the splitting step, one cluster was selected based on a pseudorandom selection weighted by the size of the clusters (number of flies best fit to the cluster). The flies are then clustered into two clusters using k-means clustering based on the expected distribution of lower level state usage. Individual HHMMs were fit to each of these new clusters. In the cluster reassignment step, we filled each unused cluster(s) with the fly(s) that had the worst fit to the current cluster assignment. After each step, we conducted 10 iterations of the EM algorithm. The sequence of perturbations was conducted for 10 iterations before a final fit using the EM algorithm was conducted until convergence.

##### 3.2. Mixture Model

To account for differing search strategies that may be utilized by different flies we also fit a mixture of HHMMs to our multi-fly data set. The goal of this model is to identify a small set of different HHMMs that can describe our entire data set by clustering flies according to the similarity in their locomotion. In this context, two flies are said to behave similarly if the same HHMM provides a good description of their behavior. To model this scenario, we added another layer to our Bayesian model of the fly movement dataset. In this topmost layer, we instantiate a Dirichlet process which probabilistically assigns a label, *z*_*n*_, to the *n*^th^ fly. When *z*_*n*_=*k*, it indicates that the *k*^th^ HHMM under consideration governs the fly’s movement.

Therefore, in addition to identifying the posterior distribution over high and low-level states we also infer a probability distribution over cluster assignments. These probability distributions are then used to determine how much we should weight each fly’s movement data when updating the parameters of the different HHMM’s.

#### 3.3 Model selection

In this study, we experimented with models with a varying number of HL states (6-16), and low-level states (4-6). One limitation of our fitting procedure is that it is not possible to compare models with a different number of states objectively, and thus the choice of model depends on the investigator. We chose a model with 10 HL states for two reasons: One reason is that different model runs with 10 HL states produced results that were more similar to each other than did model runs with either lower or higher number of states. Another reason is that as the number of HL states in the model is increased, many HL states are sparsely used. Conversely, with lower number of HL states, the distribution of observables corresponding to each HL state became broad and as a result less distinct.

### 4. Comparison of HMM with HHMM

#### Block clustering of HMM

For a given HMM fit, we may obtain the transition probability matrix (A). To obtain higher level structure from the states fit to a HMM, we may cluster the states based on their likelihood to transition to each other. One method of sorting and clustering the transition matrix into a block diagonal structure involves the use of information bottleneck formalism^9,59^. We used this method to search for a given K=10 clusters with an inverse temperature *β* from 1 to 300 and time lag values of between 1 to 45 (33 ms to 1500 ms). We found that *β* from 100-200 with time lags of 10-40 showed some of the same structure as the HHMM. We showed one solution using a time lag of 10 and a *β* of 100 on an HMM fit of 50 states with 31 used states (Figure 1-S1).

#### Bayesian model comparison

Given an HMM and an HHMM model with equal number of total states and observables O, we wish to calculate the probability of the HHMM model given the observations (P(HHMM|O)). Using Bayes theorem, we can calculate this with:

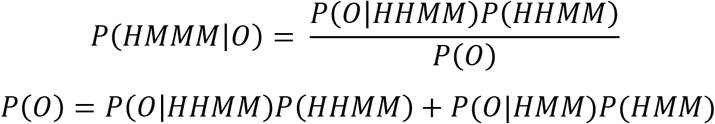

Utilizing a flat prior and Bayes factor:

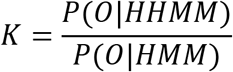

We now obtain:

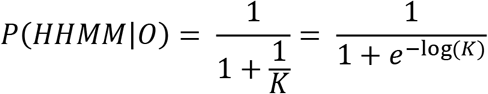

We utilized the evidence lower bound to approximate *P*(*O*|*HHMM*) and *P*(*O*|*HMM*). From this we can interpret the *P*(*HMMM*|*O*) as 1 – p-values where *H*_0_ = the HMM model is a better model fit. We obtain a *P*(*HMMM*|*O*)>0.9999 indicating that we can reject the HMM model at p<0.0001.

#### Parameter Comparison

To define an HMM, we need the transition probability p(*x*_*j*_|*x*_*i*_) for 1 ≤ *i*, *j* ≤ *K* and the description for the probability distributions defining each state. This means that there will be *K*^2^ + *K*_*n*_*p*__ parameters where *K* designates the total number of states and *n*_*p*_ designates the number of parameters defining the distributions. To define a HHMM, we need the transition probabilities at each level p(*x*_*j*_|*x*_*i*_)*l* for 1 ≤ *i*, *j* ≤ *K*_*l*_, 1 ≤ *l* ≤ *L* and the probability distributions defining each state. This means that in a model architecture in which each state at a level (*l*) is composed of equal numbers of states one level lower (*l* − 1), there will be 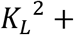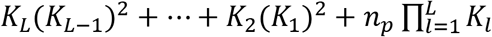 parameters. The distributions used in this report consisted of two dimensional multivariate Gaussians, which are described by 5 parameters (*μ*_1_, *μ*_2_, *σ*_1_, *σ*_2_, *ρ*). The model that we employ in the manuscript has 10^2^+10*5^2^+50*5 = 600 parameters.

### 5. Post-hoc analyses based on the HHMM model

#### Pre-processing of the model output

For each time point, the model assigns the probability that the fly is in each of the 10 HL states. We only included time instances where the model assigned a probability of >0.85 for one of the 10 states; this condition was satisfied for 81% of all data points (Figure 2B). Instances during which the flies transitioned from one HL state to another HL state and back to the same initial state in less than 5 frames (~170 ms) were reassigned to the initial HL state. Furthermore, instances during which the fly only spends one frame (~30 ms) in a certain HL state before changing states were removed. These corrections were done because such short transitions are likely to arise from noise in the observables rather than rapid transitions. Tracks of HL and LL states were extracted to compute the empirical distribution of the observables (gray dots in Figure 3A, 4 and Figure 3-S1A/B).

Each HL state track was translated to begin at the origin by subtracting by the position of the first point in the track. Then we rotated each track such that the fly is moving in the forward direction (positive y axis) at the start of the track. We considered the overall vector of the 10 frames (330 ms) before each track as the vector defining the fly’s directional intent prior to starting a track. We defined the angle of rotation as the angle between this vector and the forward direction. These translated and rotated tracks allow us to visualize the distinct types of locomotion defined by the HL states (trajectories in Figure 3C and 4).

#### Sorting of the HL states

The model output numbers the HL state in a random order with respect to the underlying distribution of observables. To better understand the structure underlying transitions between HL states, we rearranged the states from low-speed-high-turn states to high-speed-low-turn-states by sorting the HL states of the model based on the ratio of their mean speed over the variance in curvature.

#### Analysis of the effect of odors on the HL states

The nominal radius of the odor-zone defined by the radius of the odor tube was 1.25 cm^20^. However, because there was some spread of the odor outside the odor-zone, the actual radius of the odor-zone was 1.5 cm^20^. A fly was considered to be inside the odor-zone when it was within the 1.5 cm radius. Although the odor was turned on 3 minutes following the start of the recording period, because the odor was only present inside the odor-zone, we considered the first time the fly entered the odor-zone after 3 minutes (first entry) as the start of the odor period. Based on a combination of the presence of odor and the location of the fly, we parsed the data into 4 categories: These were defined as inside odor-zone before first entry (B_I_), inside odor-zone following first entry (D_I_), outside odor-zone before first entry (B_O_), and outside odor-zone following first entry (D_O_).

### 6. X-Means clustering

X-means clustering is an extension of K-means clustering. K-means clustering is an iterative algorithm that assigns data points to one of K groups based on the distance between points and the cluster centers; in most versions of K-means clustering, the number of clusters is specified by the user. X-means extends the K-means algorithm by computing the Bayesian information criterion (BIC) scores associated with a given K-means model fit, and, therefore allows a better assessment of the appropriate number of clusters in the data^60^. Following Raftery et al. ^61^. We computed the t-score based on the change in BIC given an increase in the number of clusters for N = 34 flies as follows:

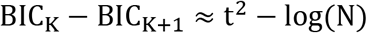

In this case, the t statistic and the corresponding P values represent the likelihood that k-means with a given cluster size will do significantly better than one with smaller cluster size. We chose the maximum cluster K that fulfilled t>3.86 (p<0.05) and within 5 percent of the minimum BIC.

In the analyses in Figure 7 and Figure 7-S3, we clustered flies based on a 10-dimensional representation of the fractional time a fly spends in each of the 10 HL states. We also employ X-means clustering in Figure 8 to cluster a fly’s response to odors. In this analysis, the clustering was performed on odor-evoked change in HL state occupancy: For inside the odor-zone, DI-BI was computed for each fly and was the input to the X-means. For outside the odor-zone, D_o_-B_o_ was computed for each fly and was the input to the X-means (Figure 8).

The clustering was performed on the 10-dimensional representation; but, for visual representation, we conducted PCA on the HL state distributions in each of the four scenarios to obtain a 2-dimensional representation of the clusters of flies (Figure 7 and Figure 7-S3). The first 2 principal components explain less than 90% of the variance.

The 10-dimensional HL state probability distributions used in Figure 7 and Figure 7-S3 reside on a 9-dimensional probability simplex:

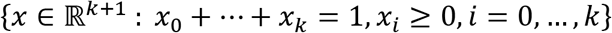

To assess whether the x-means clusters found in our data set are valid, we conducted x-means on a set of 34 points sampled randomly from the uniformly distributed 9-simplex space our data resides in. To sample from this simplex, we used a flat Dirichlet distribution marked by the following density function.

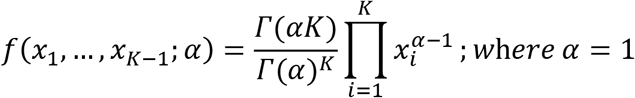

Where K = the number of dimensions. We found that x-means did not cluster these sampled data points into clusters.

### 7. Logistic Regression Model

We employed logistic regression (logit) models^62,63^–a generalized linear model which can be used to describe the relationship between independent observations and a binary dependent variable. In this study, we analyzed whether the distribution of HL states in a one-second window is predictive of whether the fly is inside the odor-zone or outside it. Because of the difference in a fly’s behavior inside and outside the odor zone, we separately performed this analysis inside and outside the odor-zone. We performed three different logistic regressions in this study: The analysis process is the same; the only difference is how flies were grouped. In the first analysis (Figure 7-S1), a <u>single</u> regression was performed for all the flies. In the second analysis (Figure 8A/B, Figure 8-S1), we calculated regressions <u>individually</u> for each fly so that there are 34 regressions–one for each fly. Finally, in Figure 8C, we calculated logistic regression for each cluster of flies–five clusters inside the odor-zone and four outside the odor-zone. The process for performing logistic regression is described below and illustrated in Figure 7-S1 and 8-S1.

First, we divide the data into 1-second time-bins. We also performed logistic regression on data subdivided into 0.33, 0.66, and 3 second bins (corresponding to 10, 20 and 90 data points) with varying amounts of time overlap and found no notable differences in model predictions.

Because we want the chance prediction to be 50% in each analysis, bins were randomly removed from either the before or during case such that the total number of bins were the same for the before period and during period. Next, we performed principal component analysis (PCA) on the distribution to obtain a smaller number of uncorrelated variables. We considered the smallest number of principal components that cumulatively explained over 90% of the variance in our analysis.

The resulting principal components were used as predictors in fitting to a logarithmic regression model. We used the MATLAB built-in function “glmfit.m” to implement fitting to a generalized linear model. For fitting to logit model, we used a binomial distribution (having experienced the odor or not) and the ‘logit’ criterion. The resulting logistic function based on the population data was used to predict if a fly was experiencing odor in any given 1-second bin.

To evaluate the predictive power of the raw observables (speed and curvature) on the behavior of the flies, the GLM was fit using the speed and curvature as predictors instead of the distributions of HL states. To compare the relative probability of correct predictions on for each fly between two different types of GLM fits (M_1_, M_2_), we considered the perpendicular distance 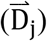 of these fits from the line of unity (indicating perfect correlation).

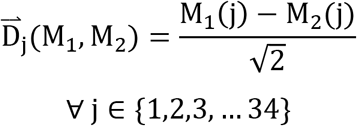

To determine whether HL states performed better than the observables, a Wilcoxon matched rank test was conducted on the 34 distances calculated for each of the model comparisons (Figure 8B, Figure 7-S1).

As X-means was conducted on the average distribution of HL states for each fly in each scenario, the partitioning obtained will not necessarily reflect the optimal clustering of flies based on ability to distinguish if the fly has encountered odor or not given a smaller (1 second) time bin observation of HLS distribution. To better partition flies into clusters based on both the average difference across time and smaller snapshots of difference across the 1 second time frame, we took X-means as the first partitioning of flies. Then we took a total of 6 flies from the largest clusters (n>10) that had the worst predictive power given GLM fits based on clusters. We then redistributed these flies into clusters in order to maximize the sum of the predictive power of individual flies across all clusters.

#### 8. Generation of synthetic flies

HHMM synthetic flies were generated based on the transition probabilities for each of the four scenarios (B_I_, B_O_, D_I_ and D_O_) separately. To generate a synthetic track, an HL state was first chosen based on the occurrence probability distribution of HL states. At this point, a new HL state was assigned for the next time instance (HL^*t*+1^) based on the HL transition probability matrix. Since the empirical flies spent variable time in the four scenarios, in the creation of the tracks we chose the median time spent. Therefore, each synthetic fly lasted until the total duration was reached for each scenario (Figure 7-S2). 100 sets of 34 synthetic flies were generated for each of the 4 scenarios (Figure 7A, Figure 7-S3). The resulting synthetic average distribution of HL states for each of the four scenarios was compared across the 100 iterations and showed high consistency between iterations.

## Acknowledgments

We would like to acknowledge the members of Bhandawat lab and Gaby Maimon for critical comments on earlier versions of the manuscript. This research was supported by NIDCD (VB), NINDS (VB) and an NSF CAREER award (VB).

## Author contributions

VB: Conceptualization; Supervision; Funding acquisition; Writing; analysis. LT: Conceptualization; Experimentation and analysis; Writing. SO: Conceptualization; Experimentation and analysis; Writing; JB: Conceptualization; analysis.

---

**Figure1-S1.**
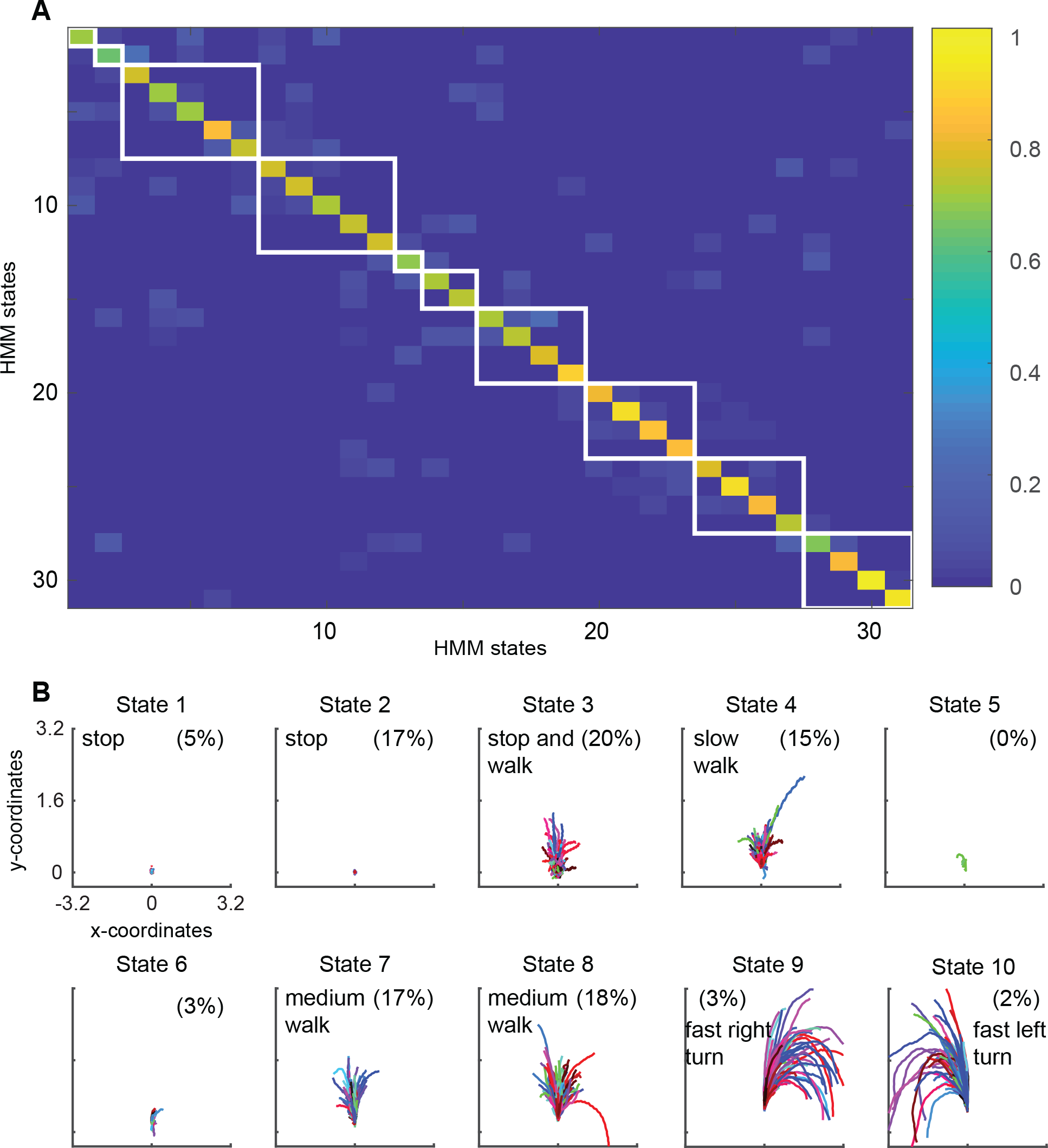
Block clustering of HMM states suggests a small number of locomotive features. **A.** Transition probability matrix of the 31 used HMM states. The states are block clustered into clusters based on the probability of transitioning to the other states. White boxes are drawn around states that are clustered together. Clustered states were sorted in the ascending order of mean speed/standard deviation of curvature, with state 1 being low-speed-high-curvature and state 10 being high speed and low curvature. **B.** Forward trajectories of the resultant clusters. States 1 through 10 correspond to the white boxes in A as shown in order from top left to bottom right.

**Figure2-S1.**
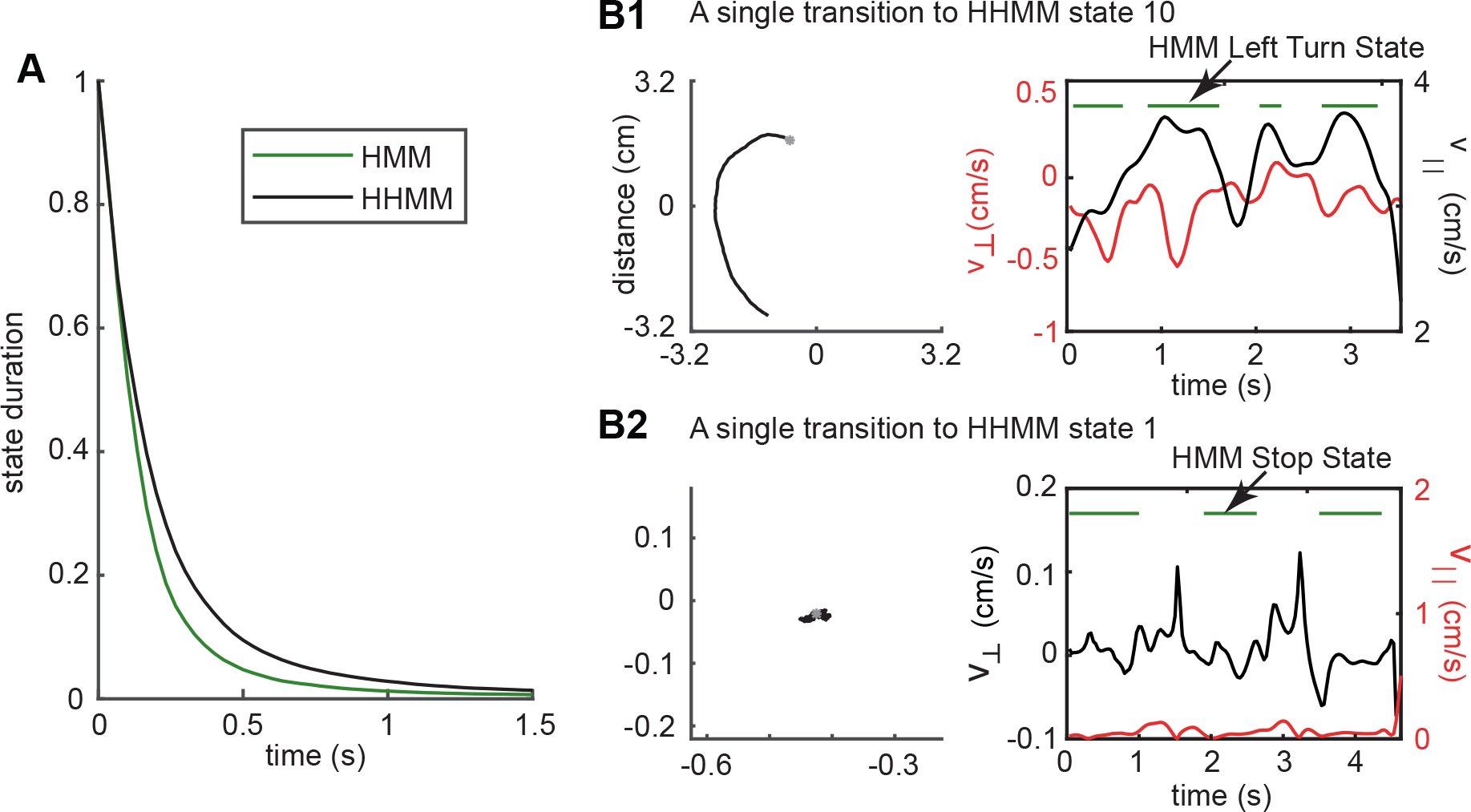
Longer duration of HHMM states allows it to discover structure in the data over longer times. **A.**Complementary cumultive distribution (cCDF) of tracks for HHMM vs. HMM states showing that HHMM states have much more tracks with longer durations. **B1**. An example track that represents a single HHMM state. HMM detects this track as a left turn but its assignment of the track as a left turn (green lines) is intermittent. **B2**. Another example - in this case for a stop state. In both examples, changes in the value of the observables result in the intermittency of the HMM state. The observable resulting in the intermittency is colored red. In the case of B1, small straight segments within the left turn result in the intermittency. In the case of B2, small movements cause an exit from the stop state.

**Figure2-S2.**
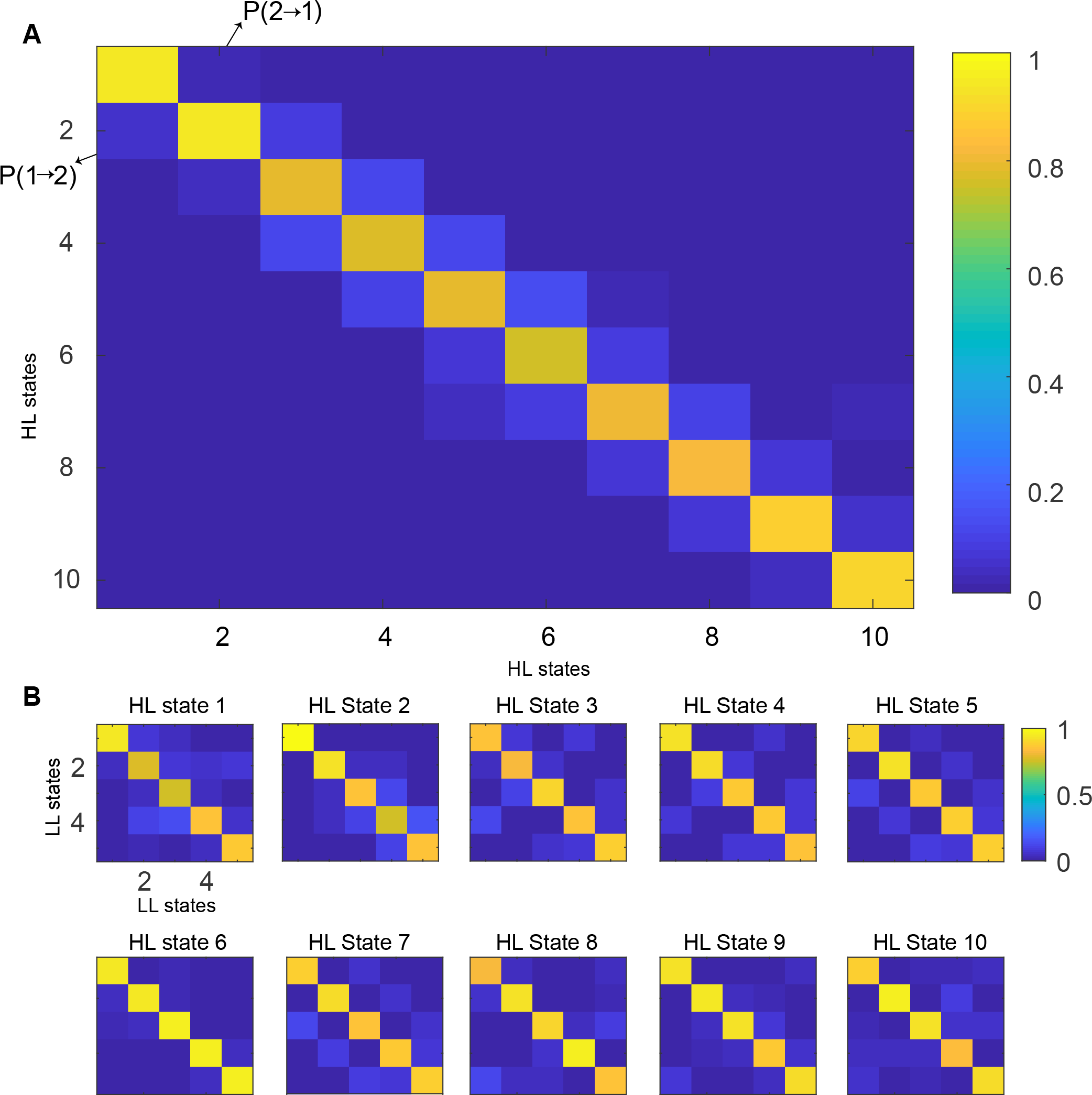
Transition probability matrices. **A.** Transition matrix for HL states. States were sorted in the ascending order of average speed/standard deviation of curvature, with state 1 being low-speed-high-curvature and state 10 being high speed and low curvature. **B.** Transition matrices of LL states for every HL state. Each HL state is described by five LL states.

**Figure3-S1.**
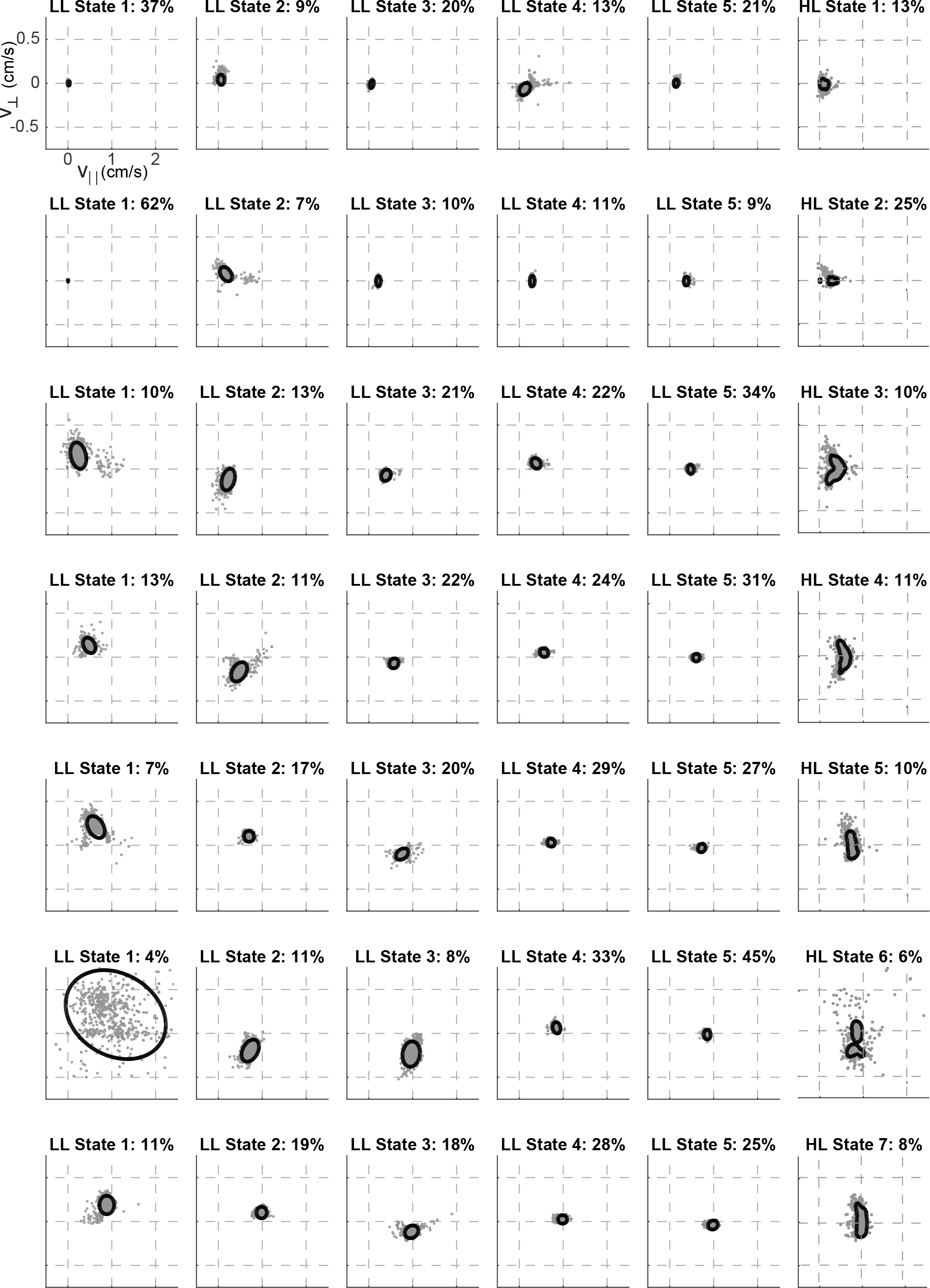
Left:HL states 1 to 10 can be computed as a weighted mixture of their LL state mvn distributions. The weghting of each LL state contribution to the HL state is based on how likely a LL state is present. Figure shows the probability of occurence of each LL state within the particular HL state. Right: Con-tour map showing modelled V_││_ and v┴ for HL states 1 to 10. HL states are computed from the weighted sum of the component LL distributions. The grey dots represent the empirical points for a particular state that are sampled from the data assigned to that state. The probability of occurence of each HL state is shown.

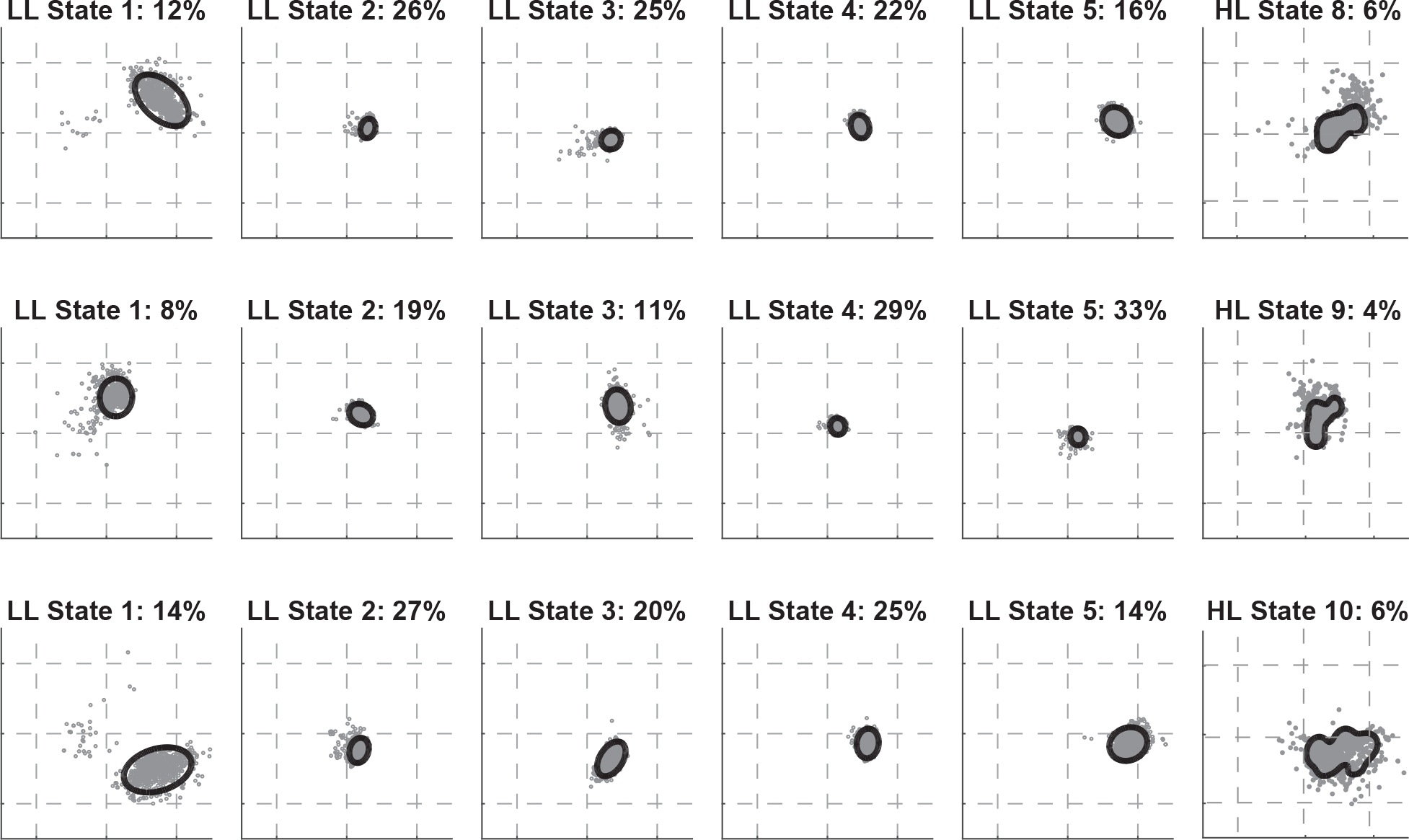

**Figure7-S1.**
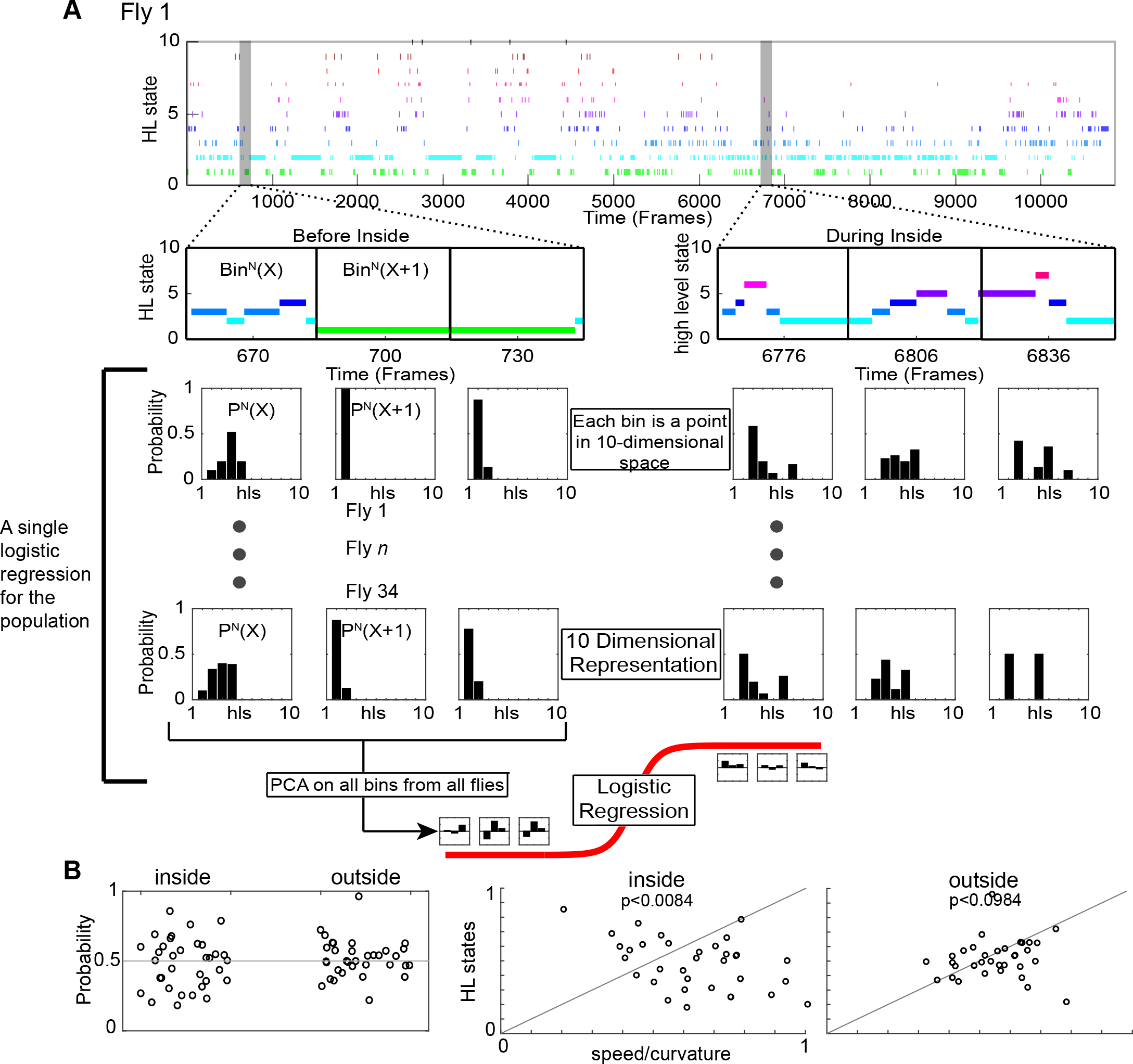
HL state distributions of the population are poor predictors for the presence of odor. We used logistic regression to evaluate whether the distribution of HL states is predictive of the presence of odor. We know from Figure 6 that ACV causes a change in the distribution of HL states, and that these changes are different inside and outside the odor-zone. Therefore, we asked whether given that fly is either inside or outside the odor-zone, can we predict based on the distribution of HL states whether the fly is experiencing an odor. In this figure, the process is schematized for inside the odor-zone (A) and the results are shown in B. More details can be found in the methods. **A.** An example of HL state as a function of time for a single fly. The HL states were segmented into 1s chunks and further subdivided into each of the 4 scenario - before inside, during inside, before outside and during outside. Here we only schematize Before Inside and During Inside because we show the logistic regression for the inside case. The HL state distributions were calculated for each segment. Thus each 1s bin is represented as a point in 10 dimensional state. We performed PCA on the entire dataset and the principal components that explains most of the variance (>90% variance) were used to fit to a logistic regression (logit) model. **B.** (Left) Results of a logrithmic regression (logit) model fit to the HL state distributions in 1 second segments for all flies do not show better probability of correct predictions over change (gray line). (Right) Distributions of high level states of the population do not improve decoding over observables.(Wilcoxon signed rank test,p<0.0084 for inside and p<0.0984 for outside).

**Figure7-S2.**
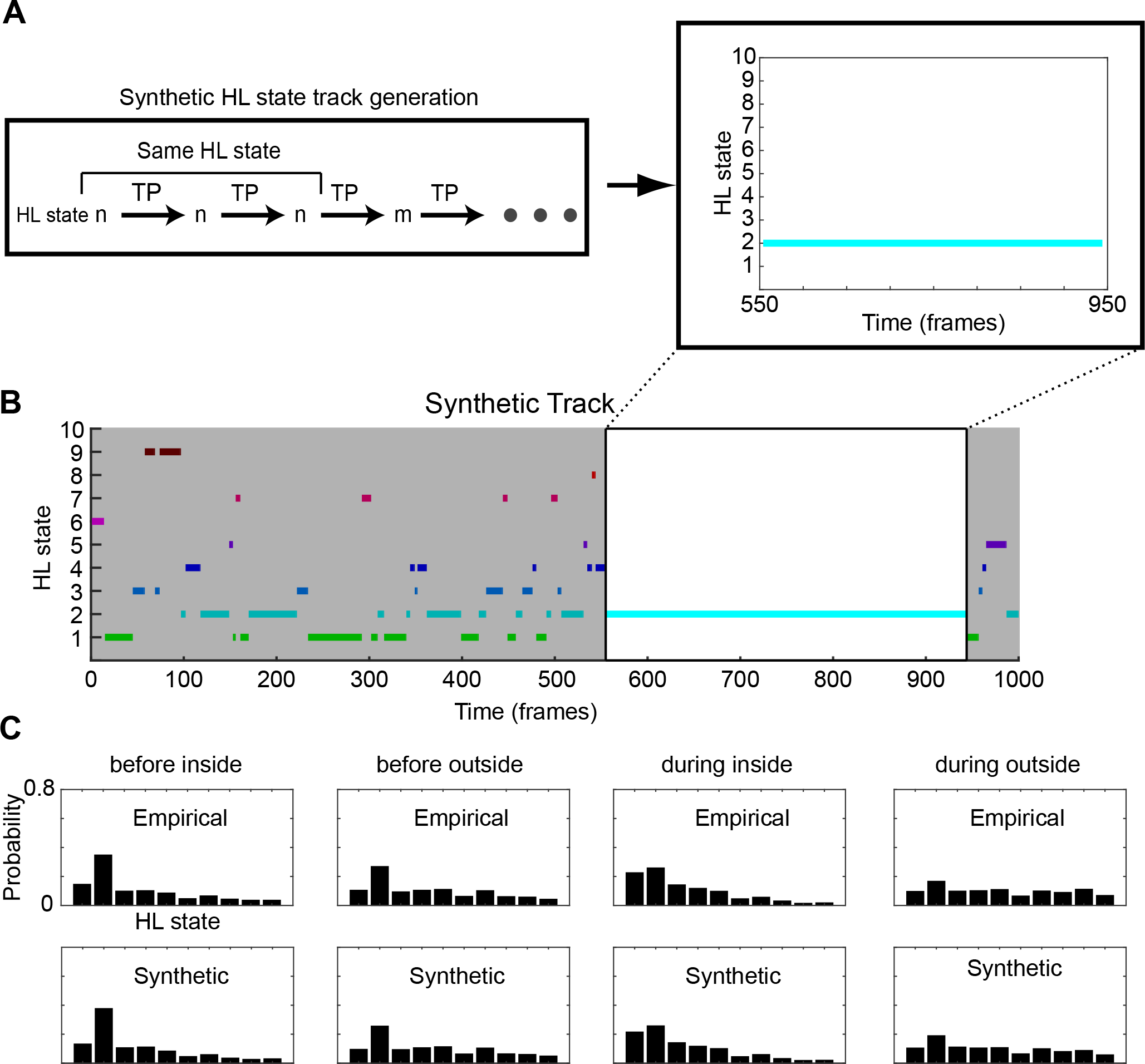
Synthetic tracks are generated for before/during and inside/outside. **A.**Flow chart for generation of synthetic flies using the HHMM model fit. A HL state is sampled from the empirical average HL state distribution for a given scenario. Progression of HL states are calculated using the global transition probability matrices associated with the HHMM and the given scenario. A fragment is shown where a synthetic fly stays in HL state 2 for 400 frames. **B.**A sample synthetic track. A synthetic track lasts for the median empirical duration of the given scenario (1000 frames are shown). **C**. The mean empirical high level state distributions are compared to the mean synthetic HL state distributions.

**Figure7-S3.**
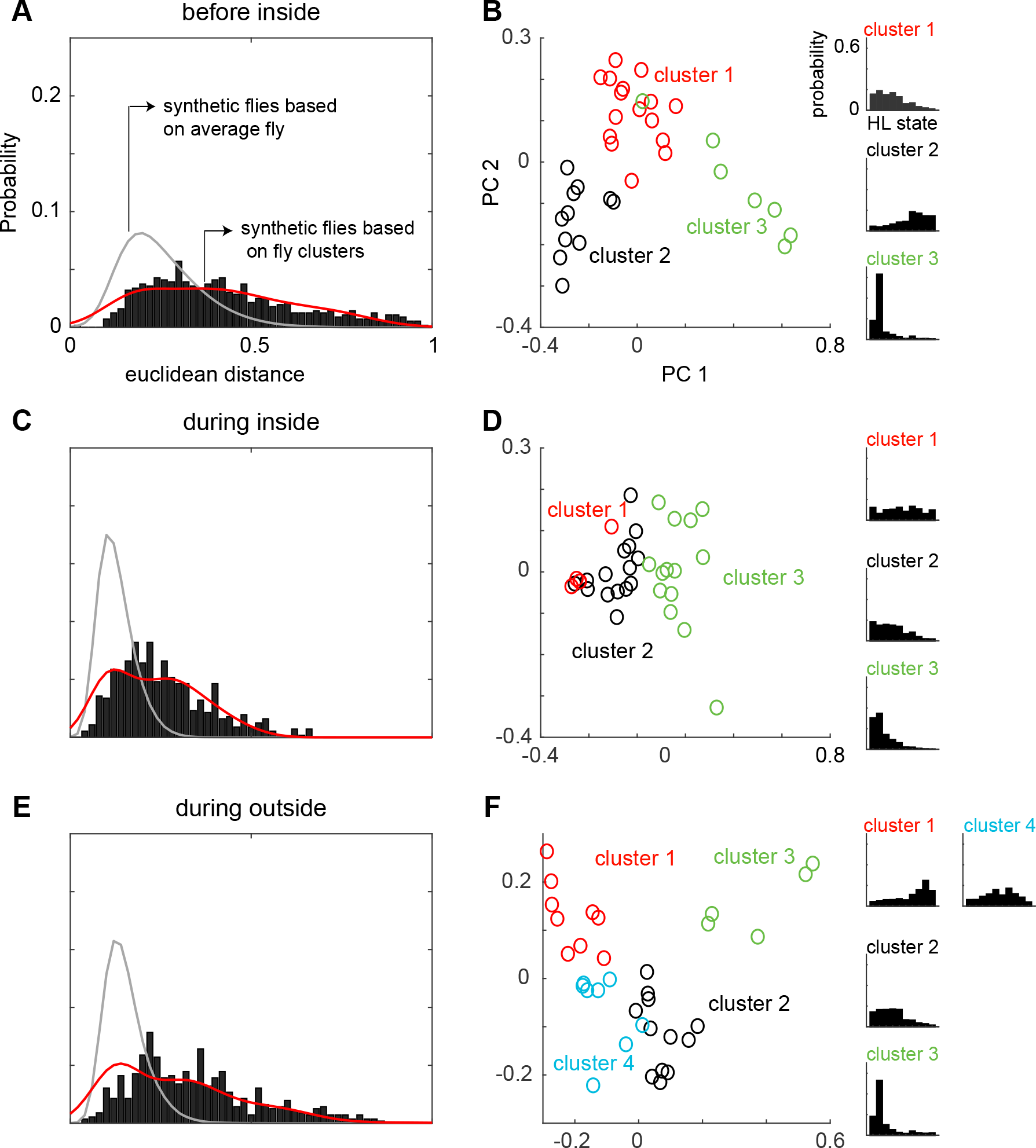
Flies can be clustered into 3-4 types based on their locomotion. **A.**The distribution of distances between flies inside the odor ring before odor. Histogram shows the empirical distances between flies in the 10-dimensional space formed by the HL states. The distance between synthetic flies based on the average fly is much smaller (gray line, Wilcoxon rank sum, p<9.57e-58). The distance between synthetic flies drawn from 3-4 clusters of flies has a distribution more similar, but still significantly different from the empirical distribution (Wilcoxon rank sum, p<4.13e-3). **B**. X-means clustering (a variant of K-means) based on the 10-dimensional space formed by the HL states show that there are 3 clusters of flies in the “before inside” section. The first two PCs are shown for visualization. Each cluster is represented by a different color. Average HL state distribution for each cluster is shown on the right. **C and D**. Same as A and B but for “during inside”. Average fly (Wilcoxon rank sum, p<3.25e-86). Cluster fly (Wilcoxon rank sum, p<0.019). **E and F**. Same as A and B but for “during outside”. Average fly (Wilcoxon rank sum, p<2.42e-130). Cluster fly (Wilcoxon rank sum, p<2.40e-10).

**Figure7-S4.**
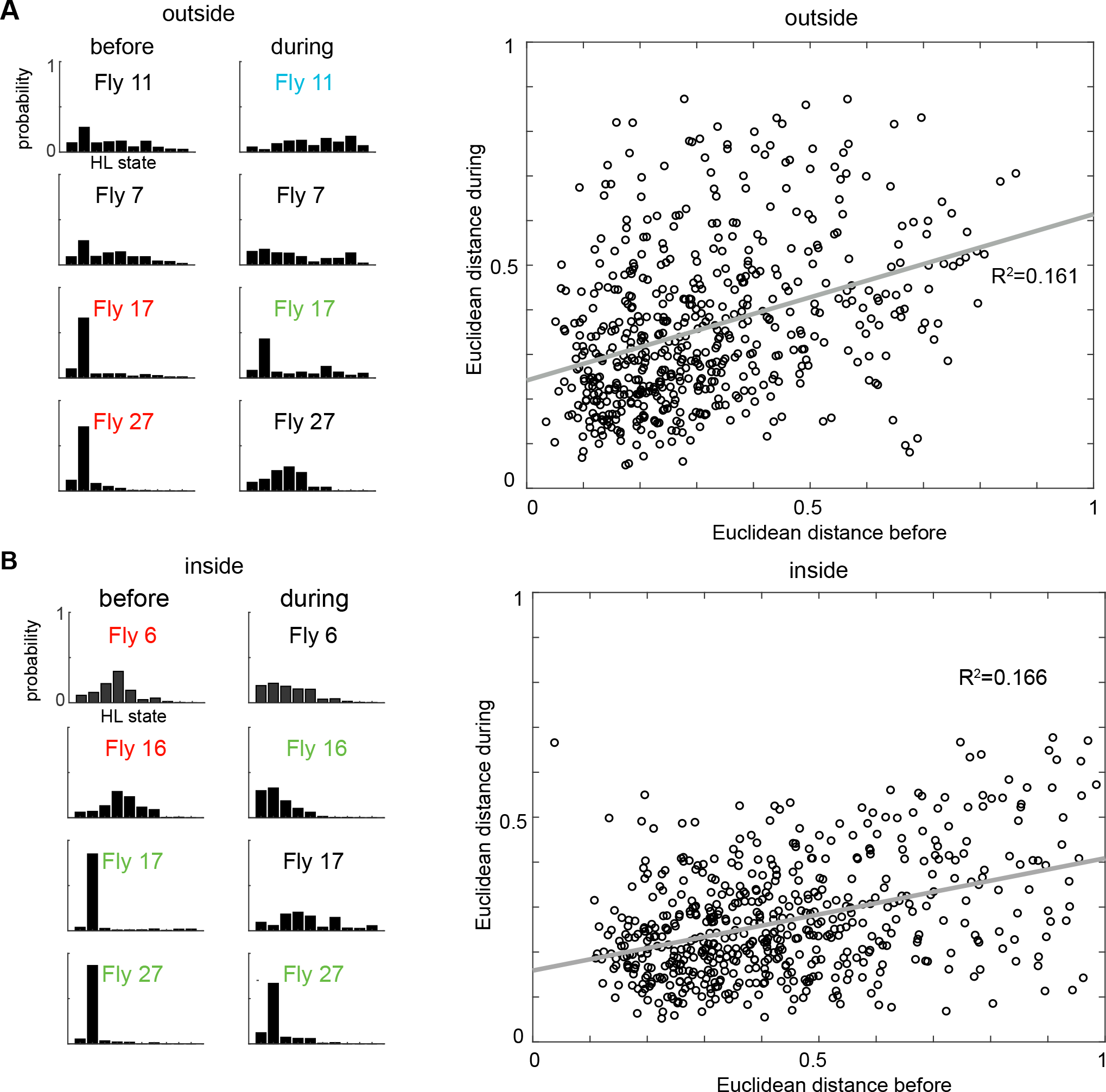
Locomotion before odor onset is only weakly predictive of locomotion during the presence of odor. **A.**Left: X-means clustering algorithm based on the average HL state distribution for each fly before the odor period and inside the odor ring grouped the 34 flies into 3 different clusters. Flies belonging to the same cluster, like 11,7 and 17,27 showed similar HL state distributions. The same algorithm grouped the flies in 3 different clusters during the odor period and inside the odor ring. The flies which belong to the same cluster before the odor period belong to different clusters following odor and show very different HL state distributions from each other. Right:Linear regression analysis between the Euclidean distances before and during the odor period outside the odor ring shows that they are weakly correlated with a r-squared value of 0.161 (p<0.001). **B.**Left: The same analysis was applied to the inside odor case. Again, flies belonging to the same cluster 6,16 and 17,27 before first entry showed different distribution of HLS usage following odor.Right: Linear regression analysis between the Euclidean distances before and during the odor period inside the odor ring shows that they are weakly correlated with a r-squared value of 0.166 (p<0.001).

**Figure8-S1.**
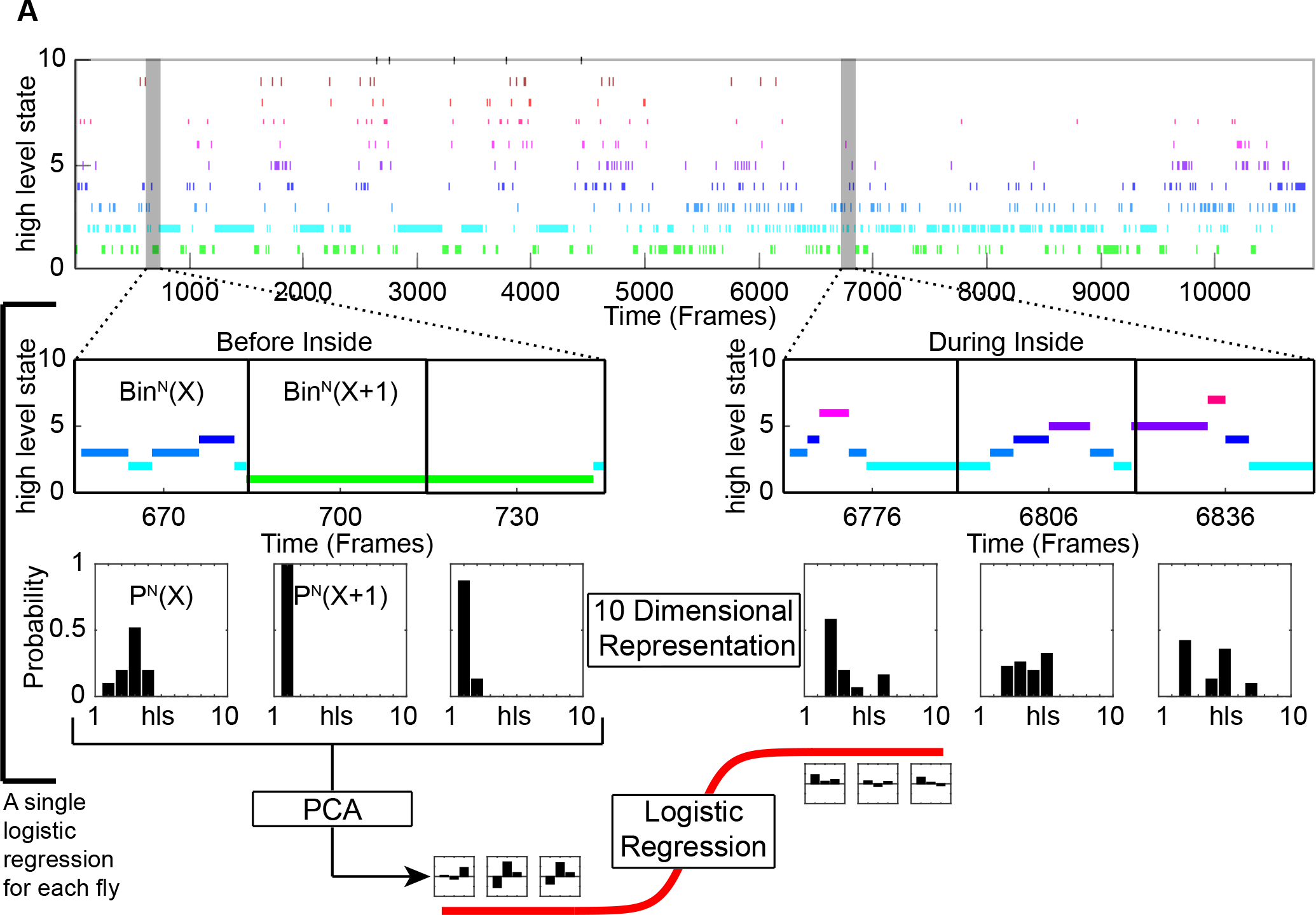
Schematic showing how logisitic regression for a single fly was performed. **A.**An example of a high level state track for a fly used in this study. The high level states were separated based on inside and outside the odor ring. For each scenario, the time series was further subdivided into before odor and following first entry. These tracks were divided into segments of 30 frames (1 second) each. The high level state distributions were calculated for each segment. PCA was applied and the pricipal components that explains most of the variance were used to fit to a logistic regression (logit) model.

